# Mimicking the Human Tympanic Membrane: the Significance of Geometry

**DOI:** 10.1101/2020.11.14.383299

**Authors:** Shivesh Anand, Thomas Stoppe, Mónica Lucena, Timo Rademakers, Marcus Neudert, Serena Danti, Lorenzo Moroni, Carlos Mota

## Abstract

The human tympanic membrane (TM) captures sound waves reaching the outer ear from the environment and transforms them into mechanical motion. The successful transmission of these acoustic vibrations in varying frequency ranges is attributed to the structural architecture of the TM. However, a limited knowledge is available on the contribution of its discrete anatomical features, which is important to fabricate functional biomimetic TM replacements. This work synergizes theoretical and experimental approaches toward understanding the significance of geometry in tissue engineered TM scaffolds. Three test designs along with a plain control were chosen to decouple some of the dominant structural attributes, such as, the radial and circumferential alignment of the collagen fibrils. *In silico* models suggested a geometrical dependency of their mechanical and acoustical responses, where the presence of radially aligned fibers was observed to have a more prominent effect compared to their circumferential counterparts. Following which, a hybrid fabrication strategy combining electrospinning and additive manufacturing was optimized to manufacture hierarchical scaffolds within the dimensions of the native TM. The experimental characterizations conducted using macro-indentation and laser Doppler vibrometry were in line with the computational models. Finally, biological studies performed with human dermal fibroblasts and human mesenchymal stromal cells, revealed a favorable influence of scaffold hierarchy on cellular alignment and subsequent collagen deposition.

Graphical abstract.
Schematic diagram illustrating the overall flowchart of the work. 3D: three-dimensional; ES: electrospinning; FDM: fused deposition modeling; TM: tympanic membrane.

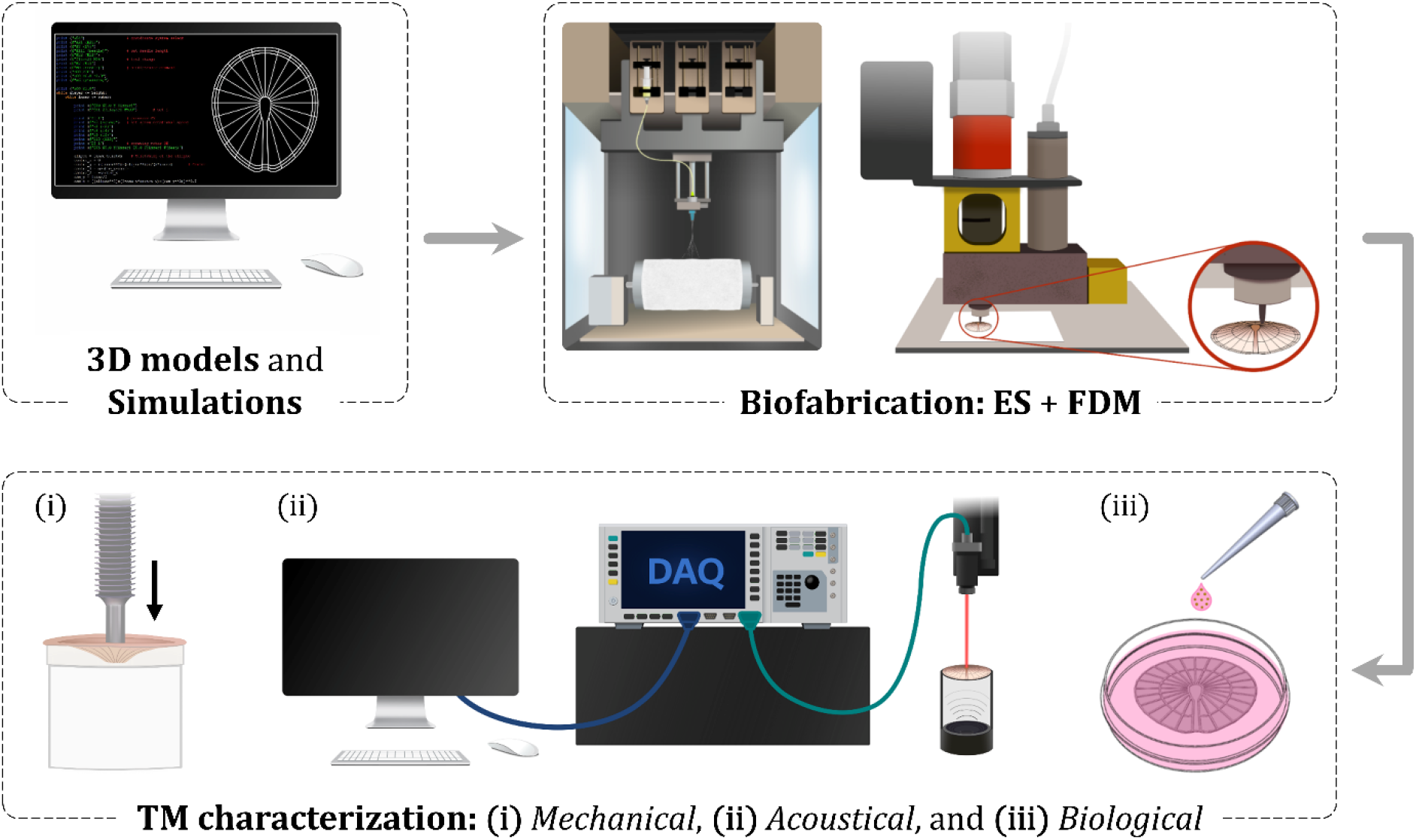

## 1. Introduction

The human tympanic membrane (TM), or eardrum, is a thin concave tissue located at the end of the outer ear canal, separating the outer from the middle ear compartments. The TM is responsible for capturing sound waves from the environment and transmitting them as a mechanical motion of the adjoining ossicular chain [1]. TM perforations are one of the most common eardrum injuries, usually resulting in a partial or severe hearing loss due to inept sound conduction. Typical etiologies inducing these perforations include acoustic traumas and microbial infections, such as chronic otitis media [2]. Microsurgical placement of autologous tissue grafts from cartilage, temporalis fascia or perichondrium is the conventional approach to repair a damaged TM. However, the incongruent tissue properties may impair an optimal hearing restoration, leading to unfavorable outcomes [3, 4]. This has often been attributed to the lack of a suitable geometry and appropriate mechano-acoustical characteristics in the recovered eardrum [5, 6].

The human TM is a tri-laminar structure, with an outer epidermal layer, a *lamina propria* consisting of fibroblasts in the middle, and a thin sheet of mucosal epithelial on the inner [7]. The concurrent sound transmission and stability of the TM is facilitated by its unique concave architecture comprising collagen fibers distributed in a radial and circumferential fashion. The distinct fiber arrangement ensures the requisite mechanical properties for transmitting acoustic waves that are converged from the outer ear through the outer auditory canal. External surface parameters of the TM, such as its area and depth of concavity have been widely studied and accurately calculated considering the interpersonal variability [8–10]. However, limited insight is available on the independent contribution of radial versus circumferential collagen fibers. In fact, with the exception of the findings reported by O’Connor *et al*. [11], where laser Doppler vibrometry (LDV) deemed the radial fibers to play an integral role in high-frequency sound conduction, this question remains largely unanswered.

The non-invasive technique of LDV has been applied extensively for characterizing the TM vibrations in middle ear diagnosis [12], and implant validations [13]. In addition, several intrusive approaches have been investigated on cadaveric eardrums for evaluating their corresponding mechanical properties [14–22]. Earlier efforts in this direction have predominantly relied on uniaxial and dynamic tensile tests; however, considering the anisotropic behavior of the TM, indentation based measurements have lately been gaining a wider acceptance [19–22]. All these studies offer a thorough characterization of the native TM, thus, guiding the development of clinically relevant tissue substitutes with comparable acousto-mechanical response.

With the recent progress in tissue engineering, novel biomaterial-based TM scaffolds have emerged as a promising alternative to the conventional tissue grafts [23–33]. Biomimetic constructs aiming to replicate the precise architecture and mechano-acoustical properties of the human TM have been investigated [23, 24]; however, the construction of a functional eardrum replacement is yet to be attained. Therefore, a better understanding of the key elements to be reproduced is critical. In this view, computational models are becoming increasingly essential to reduce preliminary empirical iterations toward design and fabrication of competent tissue constructs [34, 35]. In the past, design optimizations have often employed a combination of theoretical simulations with computer-aided design (CAD) to predict their structural response and reduce the dependence on physical scaffolds [34]. Therefore, the development of finite element analysis techniques has prompted an innovative *in silico* evaluation to validate the tissue regeneration *in vitro* and *in vivo* [36, 37].

Among the current tissue engineering strategies, biofabrication has emerged as a powerful toolbox for creating three-dimensional (3D) tissue and organ constructs [38, 39]. Several manufacturing techniques, such as, fused deposition modeling (FDM) [23, 24], 3D bioprinting [25, 26], selective laser sintering [27], and electrospinning (ES) [28–33] have been explored for the human eardrum regeneration. These technologies allow the creation of complex biomimetic scaffolds for steering cell activity, however, not many have considered the distinctive anatomy of the native TM. Among them a hybrid biofabrication strategy, combining the complementary aspects of FDM and ES, has been reported to develop TM scaffolds with precise radial and circumferential alignment [23]. An identical approach was adopted in this study for fabricating 3D hierarchical constructs within the anatomical range of the native tissue. A block copolymer of poly(ethylene oxide terephthalate) (PEOT) and poly(butylene terephthalate) (PBT) was employed as the suitable biomaterial. Coming from a family of co-poly(ether esters), PEOT/PBT has been known to exhibit desirable qualities for TM tissue engineering [23, 29].

This work aims to fill the existing knowledge gap on the significance of geometry in human eardrum. The key micro-anatomical features of the native TM were decoupled and investigated by applying relevant theoretical and experimental models. The 3D patterns chosen for this study were first analyzed computationally, following which the PEOT/PBT based TM scaffolds were biofabricated and characterized with respect to their mechanical, acoustical and biological response. A macro-indentation approach, inspired from one of the leading causes of traumatic TM perforations – foreign body instrumentation [40], was implemented to evaluate the mechanical contribution of the varying fiber arrangements, whereas, the acoustical characterization was conducted using LDV. Biological characterization of the fabricated scaffolds were performed with human dermal fibroblasts (hDFs) and human mesenchymal stromal cells (hMSCs). The cultured TM scaffolds were evaluated based on their ability to steer the cell distribution and alignment, along with an effective collagen deposition. By highlighting the influence of the chosen geometries, this work intends to demonstrate the importance of 3D patterning in TM tissue engineering.

## 2. Materials and Methods

### 2.1. Open source toolkit for generating customized TM models

A computer script was developed to design customized 3D TM scaffolds using an open source programming language – Python 3.7.0 (https://www.python.org/). The code was formulated based on eight mathematical parameters (see Supplementary Information), chosen to define the key geometrical features of the TM (**Figure 1**). These user-defined parameters facilitated the generation of a wide-range of TM models with distinct two-dimensional (2D) and 3D architectures. Mathematical algorithms were developed to convert the models into numerical control (NC) files and CAD files, respectively. Some of the essential Python packages used, include NumPy [41] and Matplotlib [42], along with the standard built-in library.

**Figure 1.**
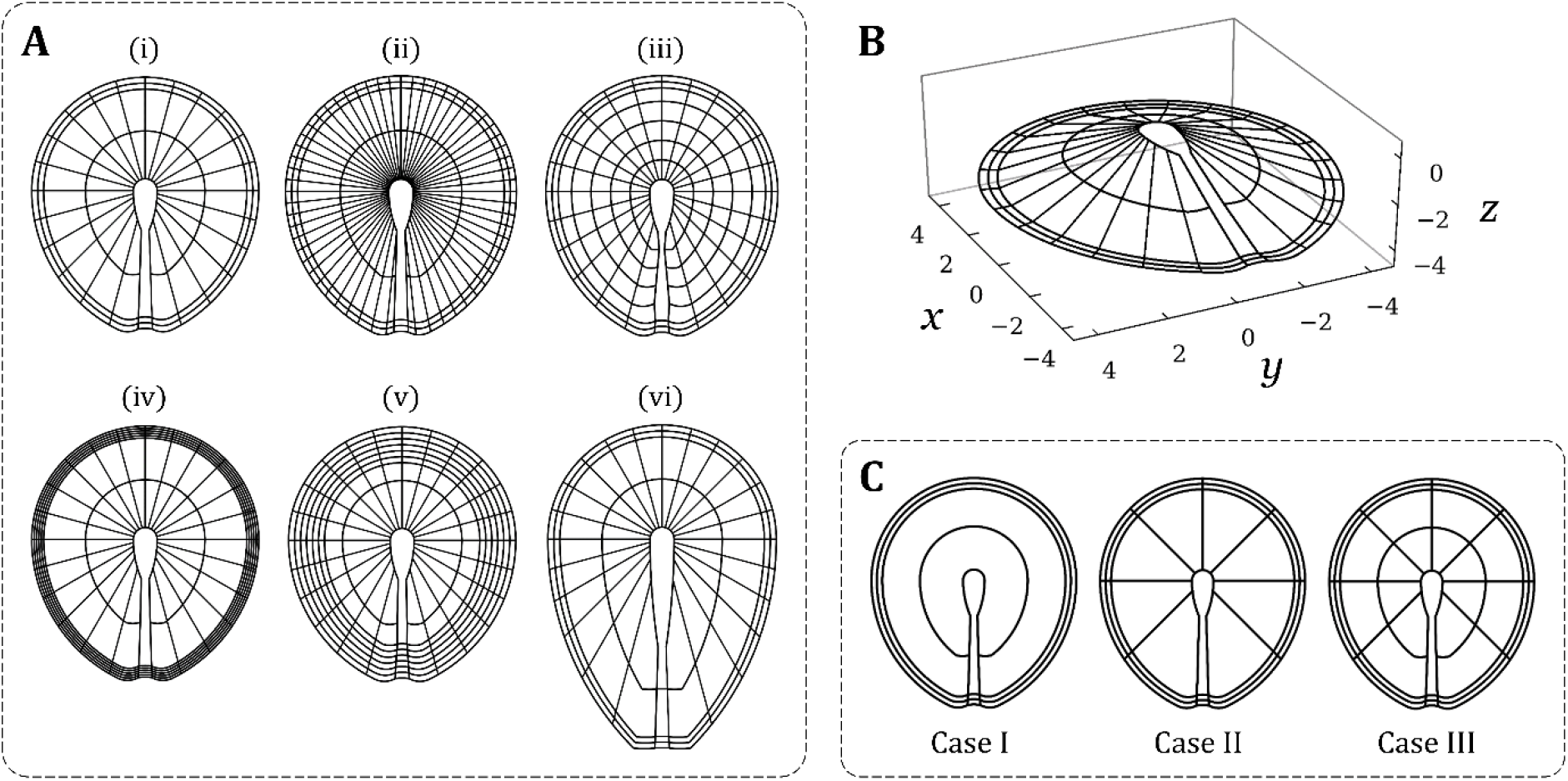
Generation of 3D TM models using a Python script. **(A)** Demonstration of the versatility of the developed script in generating structures with varying fiber arrangement, shape and dimensions **(i-vi). (B)** Conical TM models (an example shown with height 2.2 mm) can be created with the script. **(C)** Chosen test cases to decouple the key geometrical features of the native TM.

Three geometrical patterns along with a blank control were chosen as the test cases for this study. The selected designs were titled cases I, II and III, where case I consisted of only circumferential filaments, case II only radial filaments, and case III comprised the combination of both filament arrangements (**Figure 1C**).

### 2.2. Theoretical simulations: mechanical and acoustical

Computational models (COMSOL Multiphysics^®^ 5.4, Comsol B.V., the Netherlands) were designed to simulate the mechanical and acoustical behavior of the selected TM geometries. CAD files of the different FDM cases and electrospun mesh were imported and adapted as per the specifications of the physical scaffolds. The mechanical simulations were modeled based on the macro-indentation measurement (refer to section 2.4.1), using the *structural mechanics* module of COMSOL. A continuous displacement was applied at the center of the FDM filaments at a rate of 3 μm/s for 1800 s with a step-size of 0.02 s. The subsequent response of the chosen cases was obtained by evaluating their maximum normal stress and strain values, defined as the *first principal stress and strain* on the software. On the other hand, the acousto-mechanical response of these scaffolds was simulated using the *acoustics* module of COMSOL. The pre-defined multiphysics coupling of acoustic waves and structural vibrations, *acoustic-structure interaction* was configured so that the resultant models were governed by the underlying physics of both pressure acoustics and structural mechanics. The TM scaffolds were simulated within an air environment, while being subjected to a frequency sweep of planar waves, hitting them normally. An *eigenfrequency study* was performed to compute the different modes of vibrations along with their respective resonant frequencies. All theoretical investigations were carried out within the linear elastic regime of the imported geometries.

### 2.3. Biofabrication of TM scaffolds

The chosen models of the TM were constructed by applying a hybrid fabrication strategy, combining ES together with FDM. Based on prior studies in literature [23, 29], the 300PEOT55PBT45 grade of PEOT/PBT copolymer (kindly provided by PolyVation B.V., the Netherlands) was employed as the suitable biomaterial for both techniques. The selected polymer composition was synthesized by a two-step polycondensation of polyethylene glycol (PEG), where 300 represents the molecular weight (Mw, g/mol) of the starting PEG, and 55/45 constitutes the weight ratio of PEOT and PBT, respectively.

#### 2.3.1. Electrospinning

Within the dual-scale fabrication approach, ES was performed first to create a homogeneous mesh of polymeric nanofibers, replicating the collagen nanofibrils in the native TM. PEOT/PBT was dissolved in a 70:30 (v/v) solvent mixture of trichloromethane (anhydrous, Sigma-Aldrich, the Netherlands) and hexafluoro-2-propanol (analytical reagent grade, Biosolve B.V., the Netherlands), by stirring overnight at ambient conditions (**Figure S1A**). An ES setup (Fluidnatek LE-100, Bioinicia S.L., Valencia, Spain) was used to manufacture the electrospun meshes, while maintaining the environmental conditions at 23 °C and 40% of relative humidity. The polymer solution was ejected through a 0.8 mm spinneret at a flow rate of 0.9 mL/h that was kept constant while investigating the influence of other operating parameters. The overall mesh morphology and fiber diameter were optimized with respect to the polymer concentration, applied voltage, collecting substrate and air gap. PEOT/PBT concentrations from 12% to 20% (w/v) were electrospun at three different voltages – 12 kV, 20 kV and 28 kV. The effect of the air gap (10 cm, 15 cm and 20 cm) was also evaluated. All parametric characterizations were conducted for an ES duration of 5 min. A flat stainless steel collector with an aluminum foil substrate was used for the optimization studies, whereas, a polypropylene (PP) sheet mounted on a rotating mandrel (stainless steel, diameter = 200 mm, length = 300 mm) at 200 revolutions per minute (rpm) was employed as the collector for the scaffold fabrication (**Figure S1C**). The motivation behind the change in collector is discussed in section 3.3.1, along with its influence on the fiber diameter and scaffold thickness.

Scanning electron microscope (SEM; Philips/FEI XL30 Environmental SEM, the Netherlands) was used to characterize the electrospun nanofibers. Samples were gold-sputtered (Cressington Sputter Coater 108 auto, Watford, United Kingdom) for 100 s at 30 mA, and imaged at an accelerating voltage of 10 kV and working distance of 10 mm.

#### 2.3.2. Fused deposition modeling

Following the creation of electrospun meshes, 3D micro filament patterns were subsequently deposited using FDM technology. NC files with the necessary G-code commands were exported from the Python script, and loaded on to the FDM based 3D fiber deposition system (BioScaffolder, SYSENG, Germany; **Figure S2A**). The polymer loaded on a stainless steel cartridge was melted at 190 °C for 30 min, before starting to manufacturing of the patterns. Extrusion of the molten PEOT/PBT was controlled by a combination of screw-driven system and pneumatic pressure. The translational velocity was maintained constant at 157 mm/min, and a parametric optimization with respect to nozzle diameter, applied pressure, and screw rotation was conducted to obtain the smallest filament diameter possible. To achieve this, three different needle classes were investigated – (1) stainless steel with nominal inner diameter (ID) of 184 μm (ID184, BL28E03A, DL technologies, USA), (2) stainless steel with ceramic tip and ID of 100 μm (ID100, DL32005AC, DL technologies, USA), and (3) stainless steel with ceramic tip and ID of 70 μm (ID70, DL003005AC, DL technologies, USA). Subsequently, the resolution with the chosen nozzle was further optimized by testing screw speeds of 70 rpm, 50 rpm and 30 rpm, followed by applied pressures of 750 kPa, 500 kPa and 250 kPa. The resultant filament diameters were evaluated using a stereomicroscope (Nikon SMZ25, the Netherlands). Finally, the optimized FDM parameters were applied to manufacture hierarchical TM scaffolds by depositing a single layer pattern (case I, II or III) of copolymer over the electrospun scaffolds (**Figure S2B)**.

### 2.4. Characterization of TM scaffolds

The fabricated scaffolds were characterized mechanically, acoustically and biologically to gain a deeper insight into the role of geometry in the designed scaffolds for TM regeneration. The experimental setup implemented for these characterization techniques have been described in the following subsections.

#### 2.4.1. Mechanical characterization

A macro-indentation approach was developed for evaluating the mechanical response of the fabricated test cases. Inspired by the TM perforations caused by foreign body instrumentation, such as from cotton buds or other sharp objects [40], a cylindrical stainless steel probe with a diameter of 2.5 mm was used to evaluate the stress-strain profile of the developed scaffolds until the point of rupture (**Figure 4A**). The outer perimeter of the fabricated scaffolds were fixed onto the rim of a hollow tube (inner diameter = 8 mm, outer diameter = 12 mm), using a double-sided tape (Van Loenen Instruments, the Netherlands). This fixation step was performed under wet conditions to facilitate handling, followed by a drying phase of 3 hours before further testing. The dimensions of the probe and tube were chosen to ensure a uniform deformation to evaluate the contribution of all the FDM microfibers during measurement.

The uniaxial indentation was performed on a mechanical tester (ElectroForce 3200, TA Instruments, USA) with a 10 N load cell. All the test cases were evaluated (n = 4) until failure. A scan rate of 0.003 mm/s was used for the study. A Python-based analysis script was developed for the computation of relevant mechanical properties such as Young’s modulus, yield strain, resilience, and toughness. Calculations were made by neglecting the bending stresses, and assuming a membrane-like behavior of pure stretching between the central load and the outer boundary of the sample [43, 44]. The Young’s modulus (E) was calculated as the slope of the linear elastic region of the corresponding stress-strain curves. The toughness was computed by calculating the area under the stress-strain curve until the point of fracture, and the resilience was given by the area under the linear elastic region. While defining the linear range, strain values below 1% were avoided due to irregular initial contact between the scaffold and the probe. On the other hand, the upper limit was chosen until the yield strain, which is interpreted as the point of first plastic deformation. To ensure an accurate selection of the elastic regime, a linear regression was performed confirming an R-squared value > 0.95 for all the samples (**Table S1**).

#### 2.4.2. Acoustical characterization

The sound transfer function of the fabricated scaffolds was investigated using LDV. The applied test stand consisted of a 25 mm cylindrical canal, a probe microphone (ER-7C, Etymotic Research, USA) for measuring the applied sound pressure, and a laser Doppler vibrometer (sensor head CLV 700, controller CLV 1000 with modules M300, M050 and M003, Polytec GmbH, Germany) for calculating the vibration velocity. Each TM specimen was clamped at one end of the canal with a silicone coupling ring, and a multi-sinusoidal signal in the range of 100 Hz to 3 kHz at 90 dB (sound pressure level, SPL) was applied from the opposite terminal by an insert earphone (ER-2, Etymotic Research Inc., USA) to excite it. The vibration velocity generated by these excitations was integrated mathematically to compute the displacement.

The LDV measurements were performed in the center of the specimen, equivalent to the umbo at the center of the native TM. The outer FDM rings of the scaffolds were clamped onto the test stand under wet conditions with a uniform pretension. Three samples for each case were evaluated for their acousto-vibrational response and compared with that of the human cadaveric TMs. Adult human temporal bones were obtained from donors with an average age of 71.5 years (1 male, 1 female) following ethical approval from the local and national authorities. After an examination for pathological changes by an otologist, the inner ear, the incus and the malleus were removed, and the TM with the annulus fibrocartilaginous was explanted to measure its sound transfer function in the aforementioned test stand.

#### 2.4.3. Biological characterization

Two different cell types, hDFs (from dermis of neonatal skin: Lonza, CC-2509, the Netherlands) and hMSCs (from bone marrow of Donor d8011L, female, age 22: Institute for Regenerative Medicine, College of Medicine, Texas A&M Health Science Center, USA) were used independently to assess the influence of geometry on cellular alignment and collagen deposition. The fabricated scaffolds were sterilized in ethanol for 60 min, followed by an overnight drying step inside the biosafety cabinet. The sterilized scaffolds were washed twice with Dulbecco’s phosphate buffered saline (DPBS, Sigma-Aldrich, the Netherlands), and incubated for 2 hours in tissue culture media, prior to the seeding. A seeding density of 250,000 cells per scaffold was used in Dulbecco’s modified eagle medium (DMEM; high glucose, GlutaMAX™ supplement, pyruvate, Fisher Scientific, 12077549, the Netherlands) and minimum essential medium (α-MEM; GlutaMAX™ supplement, no nucleosides, Fisher Scientific, 11534466, the Netherlands) for hDFs and hMSCs, respectively.

##### 2.4.3.1. Cell distribution and alignment

The distribution and alignment of the cells seeded over the TM scaffolds was evaluated by culturing them in their respective media compositions, supplemented with 10% fetal bovine serum (FBS; Sigma-Aldrich, F7524, the Netherlands) and 1% Penicillin-streptomycin (PS; Thermo Fisher Scientific, 11548876, the Netherlands). Three different time-points (day 1, 3 and 7) were chosen for staining the actin filaments of the cultured cells using Alexa Fluor™ 488 phalloidin (Thermo Fisher Scientific, A12379, the Netherlands), and nuclei using 4,6-diamideino-2-phenylindole dihydrochloride (DAPI; Sigma-Aldrich, D9542, the Netherlands). Fluorescence images were acquired on an inverted microscope (Nikon Eclipse Ti-E, the Netherlands) at magnifications of 4x, 10x and 40x.

##### 2.4.3.2. Collagen deposition

HDFs and hMSCs were cultured in their appropriate media compositions supplemented with 1% L-ascorbic acid 2-phosphate (AsAP, Sigma-Aldrich, A8960, the Netherlands), 15% FBS and 1% PS. A higher FBS concentration along with AsAP was included in the culture media to enhance the collagen production by the cultured cells [45]. A natural collagen binding protein (CNA35) conjugated to a fluorescent dye (fluorescein isothiocyanate; FITC; kindly provided by Department of Biochemistry, Maastricht University, the Netherlands) was used to assess the extracellular collagen deposition during 7 and 14 days. High resolution images were acquired using a 25x water immersion objective (HC FLUOTAR L 25x/0.95 W 0.17 VISIR), equipped on an inverted confocal microscope (Leica SP8, the Netherlands). A White Light Laser was tuned to 488 nm (FITC), and 647 nm (AF647), whereas a separate 405 nm laser was used for the DAPI channel.

### 2.5. Image analysis

All the acquired fluorescence images were processed and analyzed on Fiji software (https://fiji.sc/). Quantification studies were performed for images captured at 10x magnification. Four distinct regions were chosen for investigating the cell distribution (**Figure S5A**) and cell alignment (**Figure S5B**). These regions were defined based on the FDM filaments of the hierarchical scaffolds, namely, (1) areas with no filament, (2) areas along radial filament, (3) areas along circumferential filament, and (4) junction areas between two filaments. For quantifying the cell distribution, intensity measurements were made in the chosen regions of interest (ROI, diameter = 200 pixels) in gray-scale phalloidin channel, which were then normalized for each specific image. The normalization was performed with a custom-built Fiji macro, developed for locating the ROI with the highest intensity in a particular image (**Figure S5C**). The extent of cell alignment was quantified using the OrientationJ plugin of Fiji, where its *measure function* was applied to obtain the coherence values in gray-scale DAPI channel [46]. Coherence of 0 implies full isotropy and that of 1 implies full anisotropy (**Figure S5D**). A bigger ROI (diameter = 500 pixels) was chosen for the orientation analysis to include the cells on the FDM filaments as well, and not just the ones aligning along them.

### 2.6. Statistical analysis

All the individual samples were assigned randomly to the different experimental groups. The number of replicates (n) and repeated experiments are indicated in the figure captions along with the statistical test performed. All data have been expressed as mean ± standard deviation. The statistical tests were performed with GraphPad Prism 8 (GraphPad Software, USA), where the statistical significances were determined by applying a one-way or two-way analysis of variance (ANOVA) followed by a Tukey’s honestly significant difference (HSD) post-hoc test (*p <0.05, **p <0.01, ***p <0.005, ****p <0.0001 and ns for p >0.05). An averaging algorithm capable of preserving the curve morphology was implemented for the analysis of LDV measurements [47]. For each measurement, the first resonance frequency was defined as the only landmark to be registered. The code provided by Ramsay *et al*. was utilized for this curve registration approach on MATLAB (R2018b, The MathWorks Inc., USA) [48].

## 3. Results

### 3.1. Design and generation of TM scaffolds

The Python toolkit facilitated the design of TM scaffolds with varying 2D and 3D macro- and micro-anatomical features. Specific parameters were defined within the script to control the overall shape, dimension, and fiber density of the radial and circumferential arranged filaments. **Figure 1A (i–vi)** demonstrates some of the illustrative 2D patterns created by the script. Taking **(i)** as the reference case, the density of radial fibers was increased in **(ii)**, and that of the inner and outer circumferential fibers in **(iii)** and **(iv)**, respectively. The inner circumferential fibers mark the actual tissue region, whereas the outer fibers were placed to ensure stability against curling and a uniform clamping during the TM characterizations. Finally, case **(vi)** highlights the versatility of the Python script in manipulating its overall shape and structure. Considering that the native TM is a conical tissue, the script also allows designing these models in 3D, with adjustable concavity. **Figure 1B** exhibits the 3D concave structure for the 2D reference case of **(i)**.

Even though the script allows modeling of intricate 3D patterns resembling the human TM, three simplified cases were chosen to decouple and understand the significance of its essential geometrical features. These have been depicted in **Figure 1C**, where case I focuses solely on the circumferential fibers, case II on radial, and case III on the combination of the two. All the subsequent investigations and characterizations were conducted on these three test cases along with a plain control.

### 3.2. Theoretical characterization of TM scaffolds

A theoretical characterization of the chosen geometries was carried out on COMSOL Multiphysics. Two independent *in silico* models were developed for simulating the mechanical and acoustical response of the selected cases. The following subsections summarize the computational results obtained from these models.

#### 3.2.1. Mechanical simulations

The mechanical simulations were performed by modeling a theoretical representation of the macro-indentation studies. The surface dimensions used for defining the boundary conditions such as prescribed displacement and rigid constraint were defined to mimic the experimental setup. For simplicity, all the domains were assumed linearly elastic. **Figure 2A** demonstrates the gradual deformation of the different cases at three arbitrarily selected time-points (T1) 0 s, (T2) 250 s, and (T3) 1000 s. First principal stress and strain values were computed over the entire range (0 – 1800 s), which were then used to calculate the resulting E. Among the control and the three chosen geometries, case III revealed the highest stiffness with E = 9.97 MPa, followed by case II with E = 9.42 MPa. The control case without any FDM filaments demonstrated the smallest E value of 2.85 MPa, whereas case I resulted in an E = 6.75 MPa, lying between the control and case II. These computations indicated an increase in stiffness with the addition of FDM patterns, where the radial fibers displayed a stronger influence over the mechanical behavior as compared to their circumferential counterparts.

**Figure 2.**
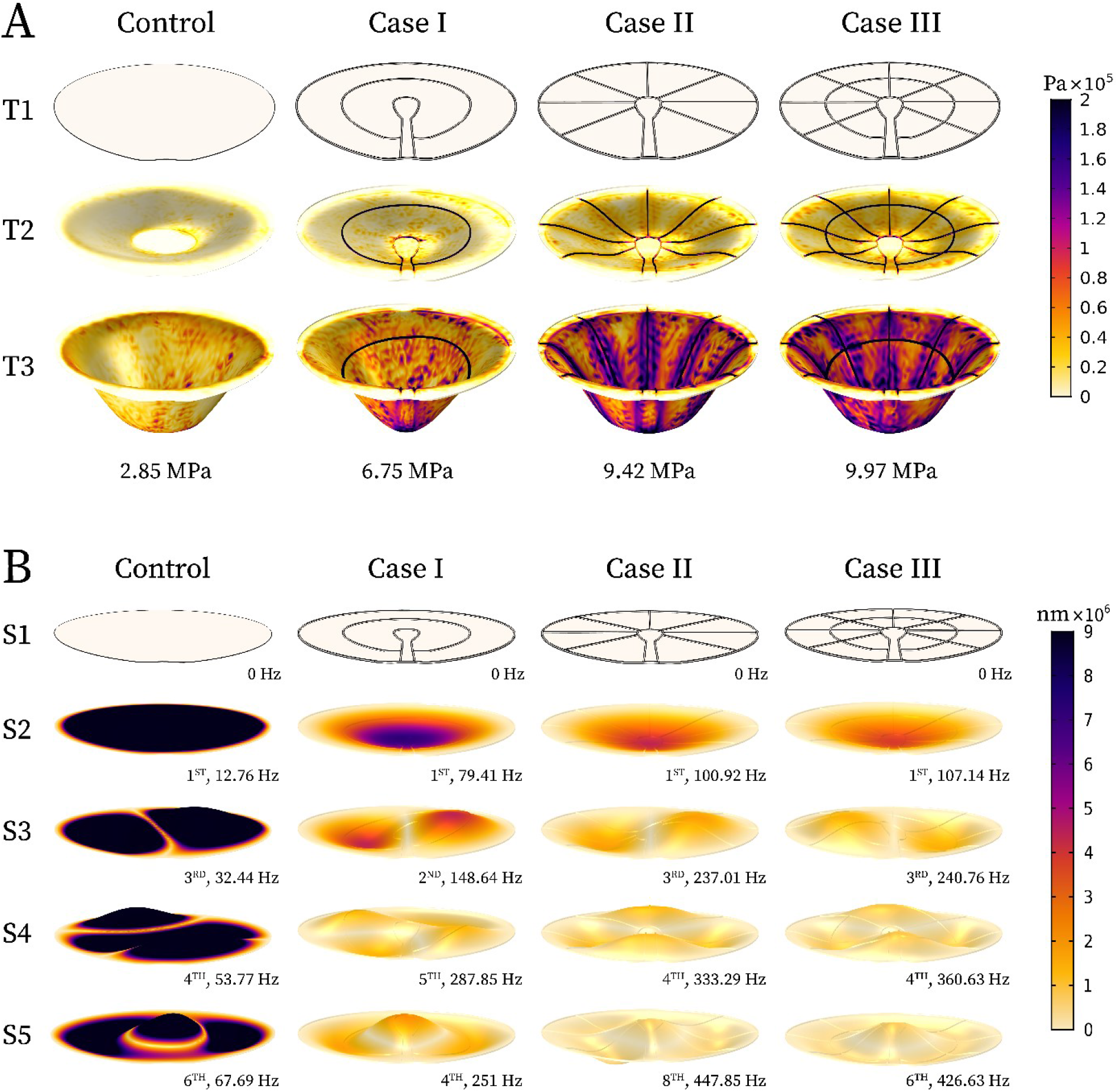
COMSOL-based computational models to predict the mechanical and acoustical behavior of the TM scaffolds. (A) Simulated mechanical response at a defined scan-rate of 3 μm/s. The scaffold deformation with time, highlighted at three different stages: (T1) 0 s, (T2) 250 s, and (T3) 1000 s. (B) Simulated acoustical response at a constant sound pressure of 0.02 Pa. (S1–S5) Key deformation shapes chosen to depict the different modes of vibrations, occurring at specific resonant frequencies of the TM models: (S1) initial state of the scaffolds, (S2) one circular node, (S3) one circular node and one nodal diameter, (S4) one circular node and two nodal diameters, and (S5) two circular nodes.

#### 3.2.2. Acoustical simulations

A computational model was developed to simulate the acoustical response of the TM scaffolds. The chosen geometries were subjected to an incident sound pressure within an air environment, to investigate its impact on their structural behavior. The corresponding natural frequencies were identified by the *eigenfrequency* analysis, which when matched the frequency of the applied wave resulted in the resonance phenomenon [49]. At their resonant frequencies, the scaffolds demonstrated extremely high amplitudes while showing characteristic vibrational shapes, known as the modes of vibration. Because of these high amplitudes, the damping was assumed to be low, and the resonant frequencies are reasoned to be very close to the natural frequencies of the scaffolds. For circular membranes, such as the human TM, these vibrational modes have been defined in terms of two key nodal points: nodal circle and nodal diameter [50].

**Figure 2B** highlights some of the resonating shapes chosen for analyzing the different test cases of this study. S1 represents the initial state of the scaffolds before any sound pressure was applied. S2 corresponds to the first mode of vibration with just one nodal circle, S3 with one nodal circle and diameter each, S4 with one nodal circle and two diameters, and S5 with two nodal circles and no diameters. The resonant frequency (RF) for each case was indicated below their images along with the mode number. In general, an increasing trend in RF was observed with rising stiffness. This was further strengthened by the fact that case II and III displayed very similar RF values, which was attributed to their comparable E values as computed by the mechanical simulations [51].

### 3.3. Optimization of biofabrication strategies

#### 3.3.1. Electrospinning

PEOT/PBT block copolymers have been widely explored for different tissue engineering applications using ES [52–55]. A couple of studies have also employed this copolymer for TM reconstruction, but lacked careful optimization of the operating parameters with respect to the native tissue [23, 29]. The key aspects investigated in this study were the electrospun fiber diameter and thickness of the collected meshes. **Figure 3A (i–iv)** demonstrates the evolution of ultrafine fiber morphology with increasing polymer concentrations at three different applied voltages. A gradual shift from large, spherical beads **(i)** at 12% (w/v) to narrow, elongated beads **(ii)** at 14% (w/v) was observed, which was followed by creation of ideal nanofibers **(iii)** at 16% (w/v) and thicker microfibers **(iv)** at 20% (w/v). Desirable mesh morphologies with consistent fiber diameters and distribution were obtained with 16% and 17% (w/v) PEOT/PBT. The influence of polymer concentration and applied voltage on the resulting fiber diameters is summarized in **Figure 3A (v)**. Furthermore, frequency distributions with a bin width of 50 nm was calculated for the fiber diameters obtained with 17% (w/v) PEOT/PBT (**Figure 3A (vi)**). Among the three voltages investigated, 12 kV demonstrated the highest convergence, thereby indicating the most homogeneous fiber distribution.

**Figure 3.**
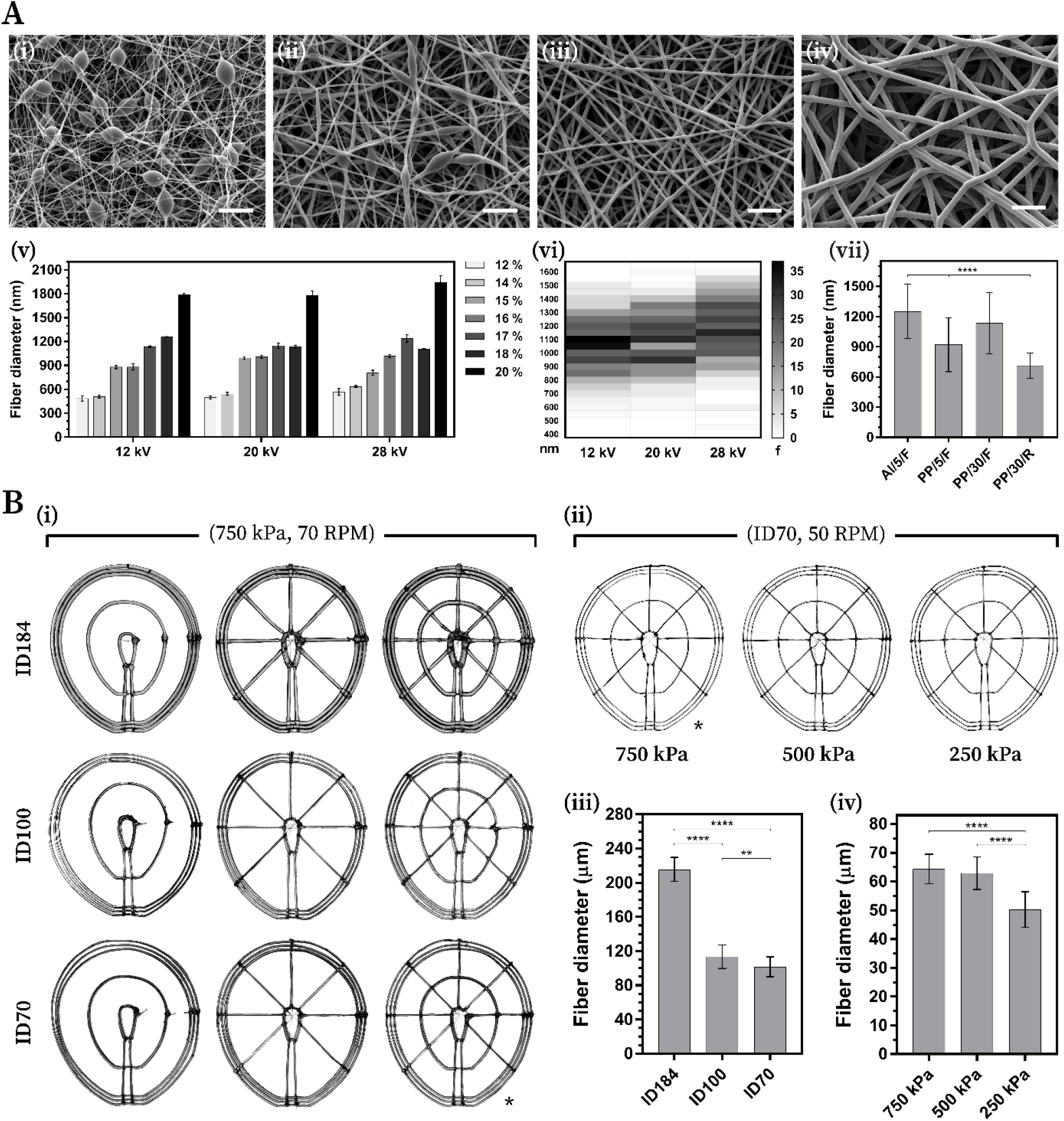
Optimization of hybrid biofabrication approach. **(A)** Optimization of the electrospinning parameters: **(i–iv)** evolution of fiber morphology from beads to nanofibers, followed by thicker microfibers (scale bar = 10 μm), **(v)** fiber diameter with respect to polymer concentration (12-20%) and applied voltage (12, 20 and 28 kV), **(vi)** heat map demonstrating the frequency distribution of the fiber diameters (left y-axis) obtained with different voltages, and **(vii)** fiber diameter with respect to collecting substrate and ES duration; conditions used for the parametric optimization: Al = aluminum foil, 5 = 5 min, and F = flat collector; conditions used for the final scaffold production: PP = polypropylene sheet, 30 = 30 min, R = rotating mandrel. **(B)** Optimization of operating parameters for the fused deposition modeling based 3D fiber deposition: **(i)** patterns produced with different needle classes (ID184, ID100 and ID70) at a constant pressure of 750 kPa and screw speed of 70 rpm, **(ii)** varying pressure for ID70 at a constant screw speed of 50 rpm; * indicates the comparison between screw speeds of 70 rpm and 50 rpm at 750 kPa with ID70, **(iii–iv)** bar graphs summarizing and comparing the average filament diameters obtained from **(i)** and **(ii)**, respectively.

Negligible differences were noted among the tested air gaps (**Figure S1D**). Considering the strong relevance of thickness in TM tissue engineering, an optimization with respect to the ES duration was conducted as well. Collection times of 20 min, 40 min and 60 min were investigated on a flat plate collector, for which a thickness of 26.69 ± 1.49 μm, 50.45 ± 0.97 μm, and 77.64 ± 2.107 μm were obtained, respectively (**Figure S1E**). Eventually, further adjustments were made in the collecting substrate to facilitate the removal of the electrospun meshes, and for increasing the manufacturing throughput. These mainly included the replacement of aluminum foil with a PP sheet, and the use of rotating cylindrical mandrel instead of a flat plate collector. The final electrospun meshes were collected on the modified setup for a duration of 30 min. The effect of these alterations on their corresponding fiber diameters is highlighted in **Figure 3A (vii)**. Overall, a statistically significant improvement was observed from an initial 1251 ± 269 nm to 712 ± 125 nm, despite the opposing trend obtained with longer collection time. A mesh thickness of 8.88 ± 0.88 μm was attained with the optimized conditions (**Table S1**).

#### 3.3.2. Fused deposition modeling

Microfilament patterns of the three test cases were deposited on top of the manufactured electrospun meshes using FDM. **Figure 3B** represents the optimization that was conducted for depositing filaments with the highest resolution possible, by obtaining the smallest filament diameter possible. **Figure 3B (i)** shows the influence of needles while printing the three test cases at a constant applied pressure and screw rotational speed of 750 kPa and 70 rpm, respectively. The smallest fiber diameter achieved for these parameters was 101.56 ± 11.29 μm with the ID70, followed by 113.40 ± 13.46 μm with ID100, which were both a significant improvement in comparison to the 215.65 ± 13.74 μm of ID184. Therefore, to further optimize the printing resolution, ID70 was investigated at lower pressures and screw rotation speeds.

A screw speed of 30 rpm with an applied pressure of 250 kPa was found to be the lowest possible combination for extruding the polymer. Although, the fabrication of scaffolds with these processing parameters were limited due to low reproducibility and difficult handling. Consequently, 50 rpm was chosen as the optimal speed, which was then evaluated for the pressure values: 750 kPa, 500 kPa and 250 kPa (**Figure 3B (ii)**). Expectedly, the highest printing resolution was attained with the lowest applied pressure, which was calculated to be 50.22 ± 6.06 μm. **Figure 3B (iii)** and **(iv)** present a summary of the fiber diameters obtained for the different tested conditions. **Figure S2C** and **Figure S2D** demonstrate the hybrid strategy for fabricating hierarchical TM scaffolds, with and without concavity.

### 3.4. Experimental characterization of TM scaffolds

#### 3.4.1. Mechanical characterization

To understand the influence of geometry over the mechanical properties of the TM, a macro-indentation system (**Figure 4A**) was developed. **Figure 4A (i)** presents snapshots taken during the measurement at four different time-points (T1 – T4). The underlying physics at each of these time-points has been explained in **Figure 4A (ii)**, where the applied load (F) results in a uniform tension (T) across the TM scaffold. A Python-based analysis was conducted to investigate the effect of this tensile stretching on the fabricated constructs. **Figure 4A (iii)** depicts the averaged stress-strain curve for each case until 60% and 10% strain. In addition, the E, yield strain, resilience, and toughness were computed for each case to present a robust mechanical characterization.

**Figure 4.**
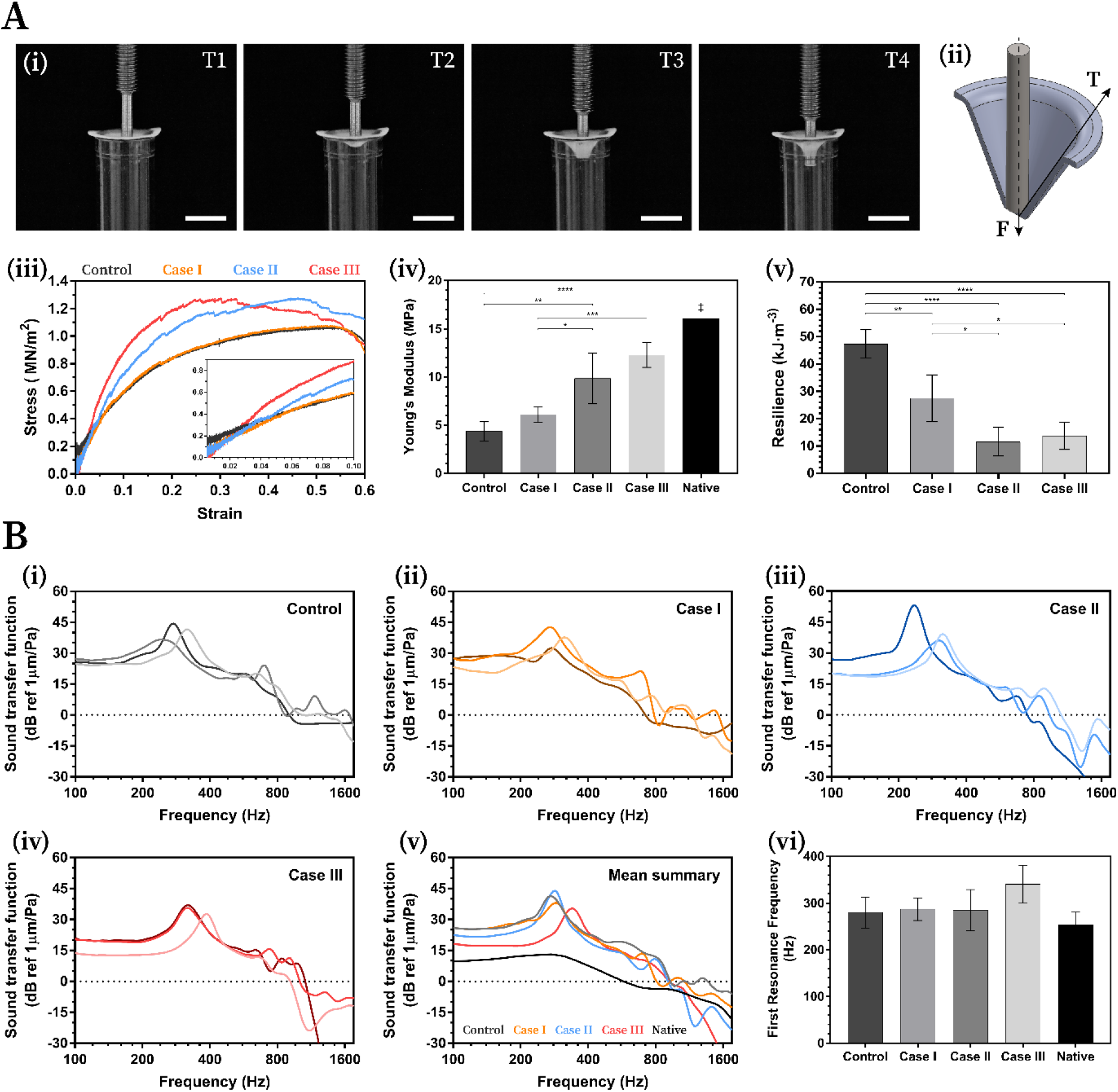
Mechanical and acoustical characterization of the fabricated scaffolds. **(A)** A macro-indentation approach was implemented for evaluating the mechanical response: **(i)** images taken during the measurement at time-points: (T1) 0 s, (T2) 250 s, (T3) 1250 s, and (T4) perforation; scale bar = 10 mm, **(ii)** schematic representation of the underlying physics, F = force applied, T = tension, **(iii)** average stress-strain curve until 60% (normal) and 10% (inset) strain, **(iv)** average Young’s moduli for the fabricated scaffolds (n = 4) compared with that of the native human TM, and **(v)** average resilience values summarized for all the cases. **(B)** Acoustic characterization of the TM scaffolds with Laser Doppler vibrometery: **(i–iv)** individual representation of datasets (n = 3) with normalized displacement plotted against the frequency sweep of the applied sound pressure wave for **(i)** control, **(ii)** case I, **(iii)** case II, and **(iv)** case III; **(v)** comparative summary of the mean curves for each case along with native TM, and **(vi)** average first resonance frequencies.

**Figure 4A (iv)** shows an upward trend in the E when moving from control (4.38 MPa) to case III (12.27 MPa). Moreover, the radial configuration of case II (9.85 MPa) resulted in stiffer scaffolds as compared to purely circumferential fibers of case I (6.1 MPa). To appreciate the clinical relevance of the obtained E values, they were compared with that of the native TM. An average E of 16.09 MPa was calculated from select literature [14, 19–21], which have been summarized in **Table 1**. Apart from the stiffness, the resilience is an important property of an elastic material. It is defined as the amount of energy that an object can absorb without creating a permanent deformation, and being capable of recovering to its initial state. **Figure 4A (v)** reveals the electrospun control to be the most resilient scaffolds followed by case I, II and III, respectively. A similar trend was also demonstrated by the yield strain (**Figure S3A**), where a statistically significant difference was observed between the samples with and without the FDM filaments. Finally, the toughness, which is interpreted as the amount of energy that an object can absorb without fracturing, was computed for all the TM scaffolds. The cases with radial filaments were noted to be comparatively tougher than the ones without them, although no statistical significance was identified. (**Figure S3B**).

**Table 1.** Reported Young’s modulus (E) values of the native human TM in selected studies: Von Békésy [14], Huang *et al*. [19], Daphalapurkar *et al*. [20], and Aernouts *et al*. [21].

| References | Technique | Sample | E (MPa) |
| --- | --- | --- | --- |
| Von Békésy, 1960 | Beam-bending | *In vivo* | 20.00 |
| Huang *et al.*, 2008 | In-plane nanoindentation | Anterior, n = 3 | 19.00 |
|  |  | Posterior, n = 3 | 17.40 |
| Daphalapurkar *et al.*, 2009 | In-plane nanoindentation | All quadrants, TM 1 | 37.80 ± 0.5 |
|  |  | All quadrants, TM 2 | 25.73 ± 2.8 |
| Aernouts *et al.*, 2012 | Microindentation | * t = 71 ± 6 µm | 4.40 |
|  |  | * t = 97 ± 16 µm | 2.10 |
|  |  | * t = 110 ± 22 µm | 2.30 |

\* t = thickness

#### 3.4.2. Acoustical characterization

The LDV technique was used to measure the velocity at the center of each sample in response to the sound excitation applied within a range of 100 Hz to 5000 Hz at about 90 dB SPL. **Figure 4B** show the calculated magnitude (normalized with respect to the applied sound pressure about 1 mm away from the specimen center) versus the frequency for n = 3 of the fabricated scaffolds and two native human TMs. The first peak in the sound transfer function denoted the first mode of vibration, corresponding to its first RF. The fundamental shape at this mode consists of one circular node at the outer perimeter of the specimen, resulting in the maximum vibration magnitude at its center.

The first RF for each measurement was located and highlighted in their respective transfer function plots (**Figure 4B (i–v)**). The average of these values were summarized in **Figure 4 (vi)** to examine the influence of the chosen geometries on the frequency response associated with the first resonance. In general, all the fabricated scaffolds demonstrated an RF value in the range of the native human TM [56]. However, the inter-sample variability was found to be high, with no clear trend detected across the investigated cases. The obtained results indicate that experimental factors, such as, the applied pretension and mass of the loaded water on each scaffold had a greater influence on its acoustical behavior as compared to the thin FDM filaments. The other modes of vibrations were not included in this study due to their diminished vibrational amplitude at higher frequency.

#### 3.4.3. Biological characterization

All TM constructs allowed an excellent growth and proliferation of the seeded hDFs and hMSCs over time. However, the different geometrical features of these scaffolds exhibited a significant influence on their cell distribution, alignment (**Figure 5**), and subsequently, collagen deposition (**Figure 6**).

**Figure 5.**
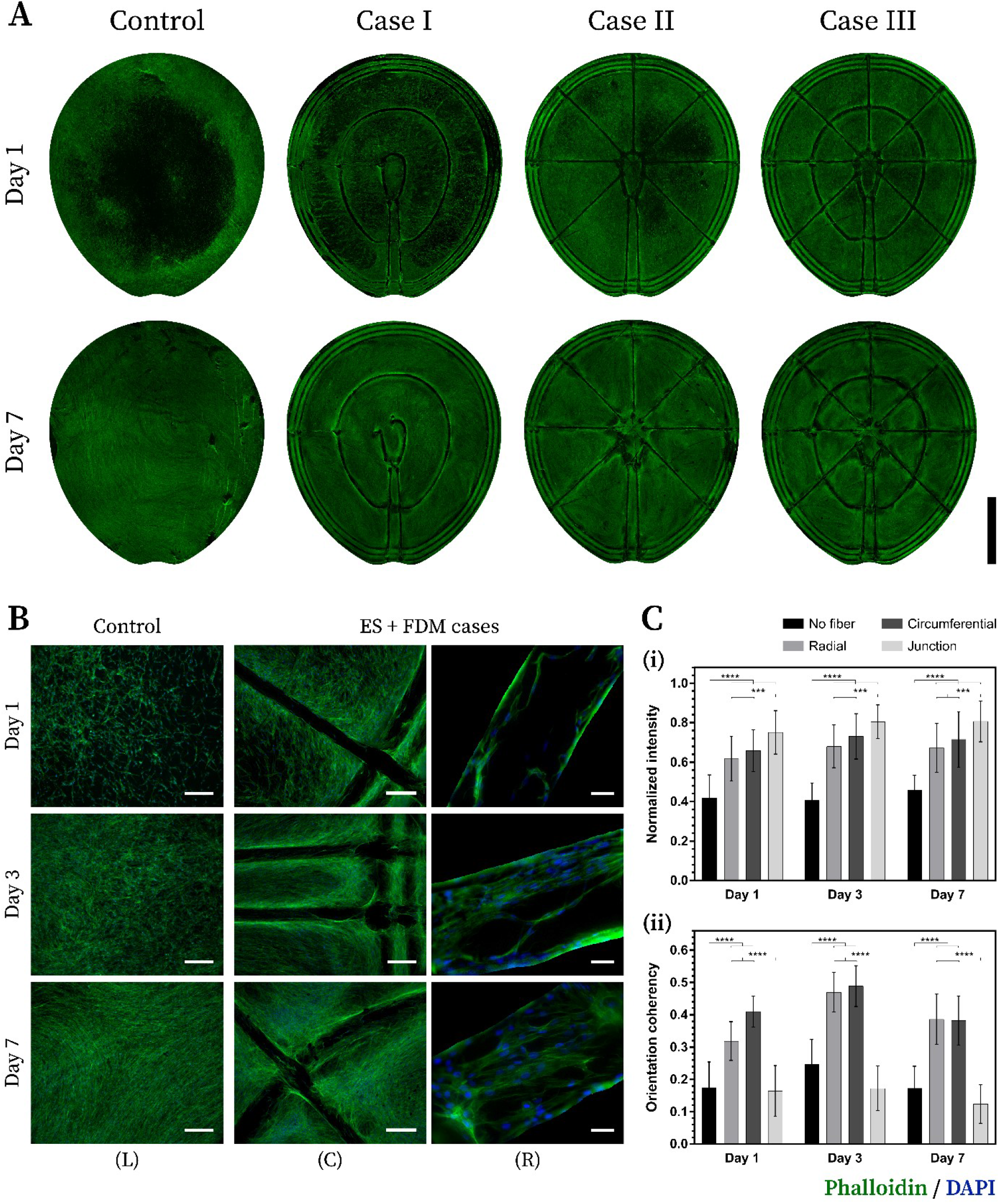
Assessment of cell distribution and alignment on the TM scaffolds. **(A)** Phalloidin staining at day 1 and 7 of human dermal fibroblasts seeded on the fabricated scaffolds; scale bar = 3 cm. **(B)** Phallodin/DAPI staining at day 1, 3 and 7 highlighting the influence of FDM filaments on guiding cell distribution and alignment; scale bar = 250 μm (lower magnification) and 50 μm (higher magnification). **(C)** Quantification of **(i)** cell distribution and **(ii)** cell alignment with respect to areas: with no filament, along radial filament, along circumferential filament, and junction between two filaments.

**Figure 6.**
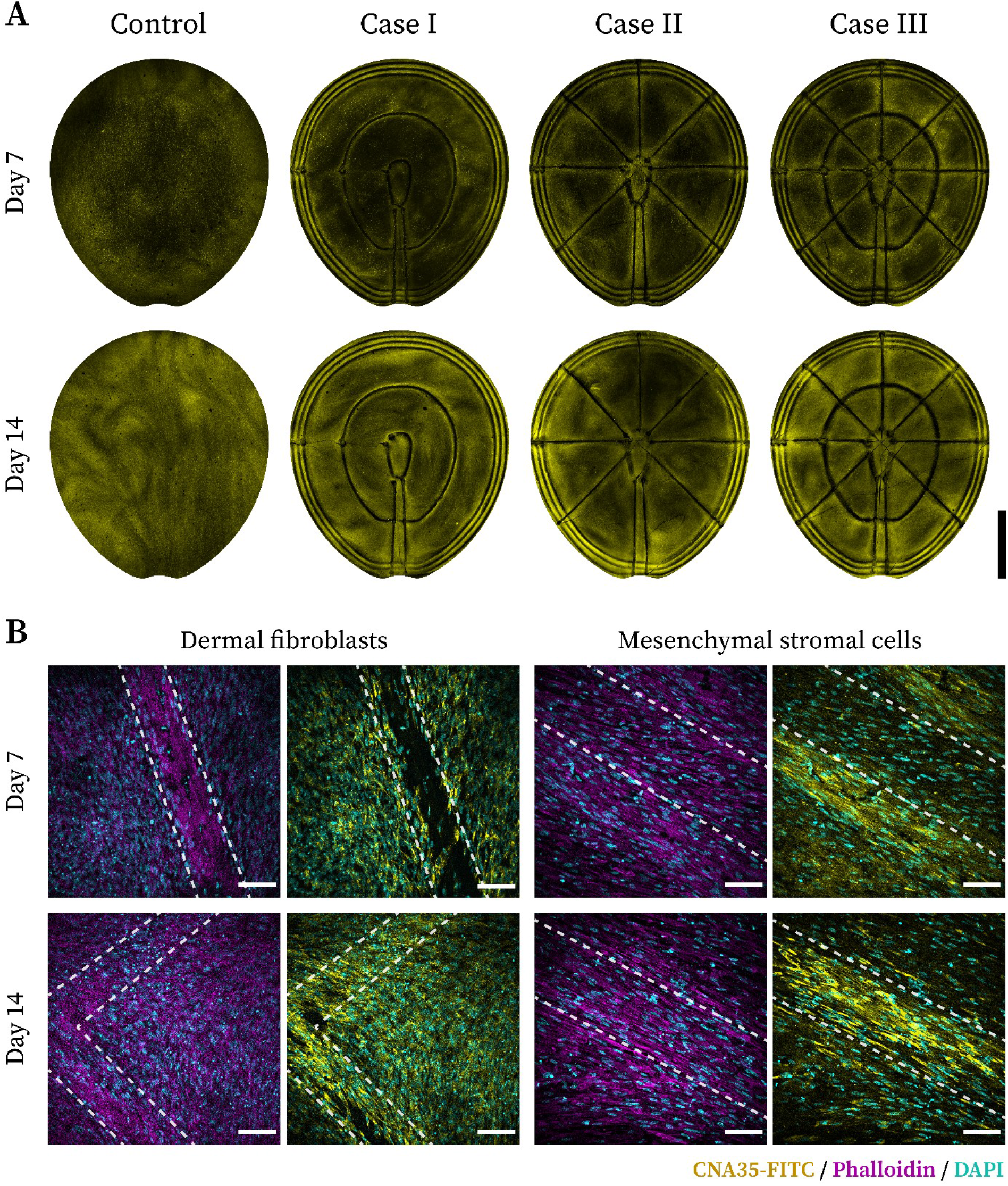
Collagen deposition by the cultured TM scaffolds. **(A)** CNA35-FITC staining at day 7 and day 14 highlights the gradual production of collagen by the seeded human dermal fibroblasts; scale bar = 3 cm. **(B)** Higher magnification images of the deposited collagen along with F-actin and nuclei: Phalloidin/DAPI (left) and CNA35-FITC/DAPI (right); scale bar = 100 μm.

##### 3.4.3.1. Cell distribution and alignment

**Figure 5A** compares the distribution of hDFs across different test cases at day 1 and 7. After 24 h in culture, a more homogeneous attachment of cells was observed in case III as compared to its counterparts. This was closely followed by case II, whereas case I and control demonstrated the worst performance in that respect. Visual inspection suggested that the presence of a higher number of closely-packed FDM filaments promoted an even attachment of cells initially. However, after 7 days in culture, all TM scaffolds were found to be completely covered with cells, indicating no major differences in terms of cell distribution.

Higher magnification images of phalloidin/DAPI stained hDFs, acquired at day 1, 3 and 7 revealed the role of hierarchical construction in steering cellular attachment and alignment (**Figure 5B**). Favorable trend with respect to cell agglomeration and orientation was observed along the FDM filaments of hybrid scaffolds (center; C) as compared to plain electrospun membranes without any FDM filaments (left; L). Moreover, the cells were noted to be aligning themselves not just along the fibers but also on top of them, as they gradually covered the entire surface (right; R).

Quantifying studies were conducted to further validate the significance of geometry in guiding these cellular responses (**Figure 5C**). Four distinct regions were chosen to analyze the hDF distribution and alignment on the fabricated scaffolds, which are, (1) electrospun mesh with no FDM fibers around, (2) electrospun mesh adjacent to radial fibers, (3) electrospun mesh adjacent to circumferential fibers, and (4) electrospun mesh at the junction between radial and circumferential fibers. The collection of cells in these specific regions were normalized with respect to each acquired image, thereby highlighting their geometrical preferences when cultured over a scaffold. A statistically consistent distribution pattern was observed over the three time-points, where junction regions demonstrated the highest agglomeration of cells, followed by circumferential and radial fibers. Areas without any FDM fibers showed the least cell density among all.

Along with guiding the cell distribution, another important aspect of hierarchical scaffolds is their ability to regulate cell alignment in the desired directions. Therefore, the orientation of the cultured cells were evaluated corresponding to the abovementioned regions. Contrary to the results of the previous quantification, the junction areas demonstrated the least alignment, which was closely followed by electrospun regions with no FDM filaments around. Cells attached along the micro-filaments, both radial and circumferential, were found to be the most aligned of all, with significant statistical differences. In general, an increment in the orientation coherency was noted for all regions from day 1 to day 3, which eventually saw a decline at day 7.

##### 3.4.3.2. Collagen deposition

After evaluating the ability of radial and circumferential fibers in steering cell attachment and alignment, a proof-of-concept study was conducted to investigate their role in guiding desired collagen deposition by the cultured cells. **Figure 6A** exhibits an increased collagen production by hDFs at day 14 as compared to day 7. This was observed across all samples; however, a careful examination revealed higher deposition of collagen around the FDM filaments, which concurs with the findings of **Figure 5**.

Higher magnification images of CNA35-FITC/phalloidin/DAPI stained hDFs and hMSCs demonstrate the growing extent of collagen, gradually enveloping the complete TM scaffolds, especially on top of the FDM filaments (**Figure 6B**). Furthermore, the deposited collagen fibrils were noted to be following the alignment of their corresponding cells.

## 4. Discussion

The effective sound conduction by the human TM has often been attributed to its unique architecture consisting of radially and circumferentially collagen fibrils. A few studies in the past have attempted to replicate similar geometrical arrangements while reporting biofabricated eardrums [23, 24, 32]. However, the majority of them have not considered it as an important aspect for mimicking the mechanical, acoustical and biological properties of the native tissue [25, 26, 28–31, 33]. Interestingly, even the ones that did focus on recreating the appropriate anatomy, lacked consistency in their chosen geometries. For instance, Seonwoo *et al*. highlighted only the radial arrangement as a crucial element for fabricating functional TM scaffolds [32]. Therefore, this work aimed to expand the current knowledge available on the role of scaffold geometry to guide the future biofabrication strategies for TM regeneration.

The Python script developed enables researchers to generate a wide range of TM architectures, which could be adjusted and implemented according to study specific objectives. In the context of the present study, three simplified geometries were chosen to decouple the key micro-anatomical features mimicking the human eardrum. The selected cases along with a plain control were first investigated theoretically, following which, a thorough experimental characterization was conducted to evaluate their mechanical, acoustical and biological responses. The computational simulations indicated a stronger influence of radial fibers on the scaffold properties than circumferential. A gradual increment in the E and RF were noted across the different geometries, with case II and III displaying the maximum values. A higher E implies a stiffer scaffold, which is also confirmed by higher RF for those specific cases. Under ideal conditions with negligible damping, the RF of a system is directly proportional to its stiffness (k) and inversely proportional to its mass (m), which is represented by 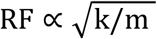 [51]. Therefore, both mechanical and acoustical simulations highlight the significance of geometry in modulating the intrinsic scaffold properties. Moreover, these models further demonstrate that by applying a suitable combination of radial and circumferential fibers, TM scaffolds with desirable acousto-mechanical response can be designed.

Experimental analyses were conducted on PEOT/PBT scaffolds, fabricated by optimizing a hybrid strategy involving ES and FDM. The microphase separated soft, hydrophilic PEOT and the hard, semi-crystalline PBT segments of the copolymer, allow a wide range of mechanical properties, thermal stability, tunable degradability, and surface energy [57]. Among them, the 300PEOT55PBT45 grade has been reported as a promising composition for the TM reconstruction [23, 29]. During the process of ES, as the applied voltage is increased, the electric field exerts a pulling force on the polymer droplet, countering the existing surface tension. As a certain threshold voltage is reached, the droplet approaches the shape of a cone, known as the Taylor’s cone and emits a jet of polymer [58]. The cone-jet during the stretching and whipping phase of 17% (w/v) PEOT/PBT was spotted around 12.5 kV, which corroborates the findings of the fiber distribution. Therefore, 17% (w/v) at 12.5 kV were chosen as the optimal parameters on a rotating PP sheet. The resultant fiber diameter of 712 ± 125 nm is a significant improvement (2.5-fold reduction) compared to the available literature on using electrospun PEOT/PBT scaffolds for TM tissue engineering [23, 29].

Subsequently, hierarchical constructs were manufactured by applying the optimized FDM parameters. The 3D deposition of filaments with a diameter of 50.22 ± 6.06 μm is a four-fold increment with respect to the previously reported resolution for PEOT/PBT copolymer [23, 59, 60]. The human eardrum has been calculated to have a diameter of 7-8 mm with a thickness of roughly 70 μm [19]. Therefore, the implemented fabrication strategy allowed the creation of radially and circumferentially patterned TM scaffolds within the anatomical dimensions of the native tissue, which is a notable achievement in comparison to the past attempts in this direction [23, 24]. Furthermore, efforts were made to construct these TM scaffolds with a concavity, which will be further investigated in future studies.

While characterizing the fabricated constructs, a macroscopic indentation was preferred over the more conventional micro or nano approaches for TM [19–21]. The macro-indentation setup assessed the contribution of the overall scaffold geometry and laid out clear distinctions between the chosen test cases. Objects subjected to central point–load deflections often experience a combination of bending and stretching stresses [43]. For instance, the behavior of a circular disc transitions from pure bending in plates to that of pure stretching in membranes, depending on the applied load and sample dimensions. The bending effect has been reported to be significant only if the resultant deformation is small with respect to the disc thickness [44]. Therefore, considering the thin anatomy of the TM scaffolds as compared to the large displacement applied, the bending stress experienced by them was disregarded in the current investigation. The stress-strain curves were evaluated by approximating a tensile stretching between the indenting probe and the outer perimeter of TM scaffolds. As predicted by the theoretical simulations, cases with the radial fibers demonstrated a similar mechanical response as compared to the rest. A gradual increment in the E value was noted from control to case III, which in the end, was found to be the closest to native TM. An average E value for the human eardrum was calculated by selecting relevant publications with comparable test setups [14, 19–21]. Furthermore, the constructs also showed promising results when examined in terms of their resilience and toughness. The native TM has been reported to rupture at overpressures as low as 35 kJ.m^−3^, with half of them projected to be damaged by 104 kJ.m^−3^, and all of them by 244 kJ.m^−3^ [61, 62]. Therefore, in general, a higher resilience is desirable to diminish the chances of a permanent deformation due to traumatic injuries. Among the tested cases, the circumferential pattern outperformed its radial counterpart. Finally, all the TM scaffolds displayed a toughness greater than 412 kJ.m^−3^, which was judged adequate as per the above recorded limits.

Moreover, the LDV measurements revealed the fabricated scaffolds to be acoustically comparable to the human TM. A similar first RF was noted across all the samples. However, a higher damping was observed in case of the native tissue, which led to a flatter peak with respect to the others. From a sound transfer standpoint, the relatively elevated magnitudes of TM scaffolds could be advantageous in compensating for the additional load due to cultured cells, but it remains yet to be investigated. No clear distinction was detected among the chosen architectures, which could be attributed to two key factors: (1) application of low pretension, and (2) the thin FDM filaments might have a negligible impact as compared to the water loading capacity of the porous electrospun network. Furthermore, with the experimental error highlighted across multiple samples (n = 3) of the same condition, the variation in the presented geometries were found to be insignificant to make any conclusive comparison.

Steering cell-matrix interactions to generate desired cellular responses have been one of the fundamental goals of biofabrication [63]. The biological characterization serves as a proof-of-concept demonstrating that the geometry of a TM scaffold has a direct impact on its tissue regeneration abilities. Two key aspects were investigated in this respect: (1) cell distribution and alignment after seeding, and (2) deposition of extracellular collagen after 14 days in culture. The hDFs were chosen as the primary cell type in this study considering their phenotypic similarity to the native cell population in the TM. The obtained results were corroborated by a parallel investigation conducted with the hMSCs. The stromal cells have been applied extensively for the TM regeneration [64], where recently, Moscato *et al*. reported their successful differentiation toward the TM fibroblasts [33].

Numerous fabrication strategies have been implemented in the past for a controlled 3D positioning of cells [65, 66]. The presence of FDM filaments in this work was found to exhibit a similar role in the homogenous attachment of hDFs on the TM scaffolds. Moreover, a quantification of the acquired images revealed a higher cell agglomeration around the filaments, which could be attributed to the presence of additional cell anchorage sites as compared to the regions devoid of any hierarchy [67]. Along with guiding the cell distribution, the hybrid scaffolds also demonstrated significant control over the cell alignment. However, it is worth mentioning here that even though the junction regions carried the highest cell density, they presented the lowest orientation coherency due to counteracting directions of the radially and circumferentially aligned cells. Therefore, appropriate adjustment should be made in the TM designs to eliminate intersecting fibers by introducing programmed discontinuities. Most of the current biofabrication approaches for human eardrum aim at developing a biomimetic platform where the strategically placed cells can deposit their own extracellular collagen matrix (ECCM), meanwhile the polymeric scaffold degrades. Several studies have investigated the impact of topographic cues on the spatial organization of the synthesized collagen [68, 69]. This work explored the role of different TM-specific geometries in steering the ECCM production and alignment. Phalloidin staining indicates that by day 7, all the cultured scaffolds were completely covered by cells; however, the extracellular collagen was spotted only along the FDM filaments. At day 14, an overall increment in the collagen deposition was noted, with sharp differences in their orientation pattern across the tested cases. Future work in this direction will focus on the expression of targeted collagen types, specific to the native TM, while initiating a simultaneous degradation of the polymeric framework.

## 5. Conclusion

This work presents a combination of theoretical and experimental approaches toward understanding the significance of scaffold geometry for TM tissue engineering. A previously reported biofabrication strategy was applied and optimized to manufacture TM scaffolds within the dimensions of the native tissue. The programmed generation of 3D designs and architectures demonstrated in this study offers a promising perspective on fabricating patient-specific tissue constructs. However, relatively simpler patterns were adopted to decouple the key structural attributes of the TM, namely radial and circumferential collagen fiber alignment. Preliminary computational modeling of the chosen cases suggested a geometrical dependency on their mechanical and acoustical response. Furthermore, the presence of radially-aligned fibers was observed to have a more prominent effect over these properties as compared to their circumferential counterparts. The experimental characterizations using macro-indentation corroborated this finding by identifying that scaffolds with a radial alignment showed E values closer to the human eardrum, whereas the circumferential fibers were deemed critical for maintaining an optimal resilience and structural integrity of the fabricated constructs. In addition, the LDV measurements demonstrated that all the fabricated scaffolds to be acoustically comparable to the native TM. Biological studies performed with hDFs and hMSCs, revealed a favorable influence of 3D hierarchy on cellular alignment and subsequent collagen deposition. Therefore, this study concludes that a combination of these micro-anatomical features is essential for the manufacturing of functional TM scaffolds. Desirable mechanical and acoustical behavior, analogous to that of the native membrane, can be achieved solely by manipulating the geometrical design of any TM scaffold. Future work in this direction will be focused on applying this knowledge to design relevant 3D architectures for TM reconstruction.

## 6. Acknowledgements

The authors declare that there is no conflict of interest. This study was supported by the *4NanoEARDRM* project, funded under the frame of EuroNanoMed III. Parts of this research have been funded by the German Federal Ministry of Education and Research, FKZ: 13XP5061B. The authors would like to thank Mr. Pratik Bachhav (BITS Pilani, India) and Dr. Ravi Sinha (Maastricht University, the Netherlands) for their assistance with graphic designing and mechanical characterization, respectively.

