## Supplementary Information for "Mimicking the Human Tympanic Membrane: the Significance of Geometry"

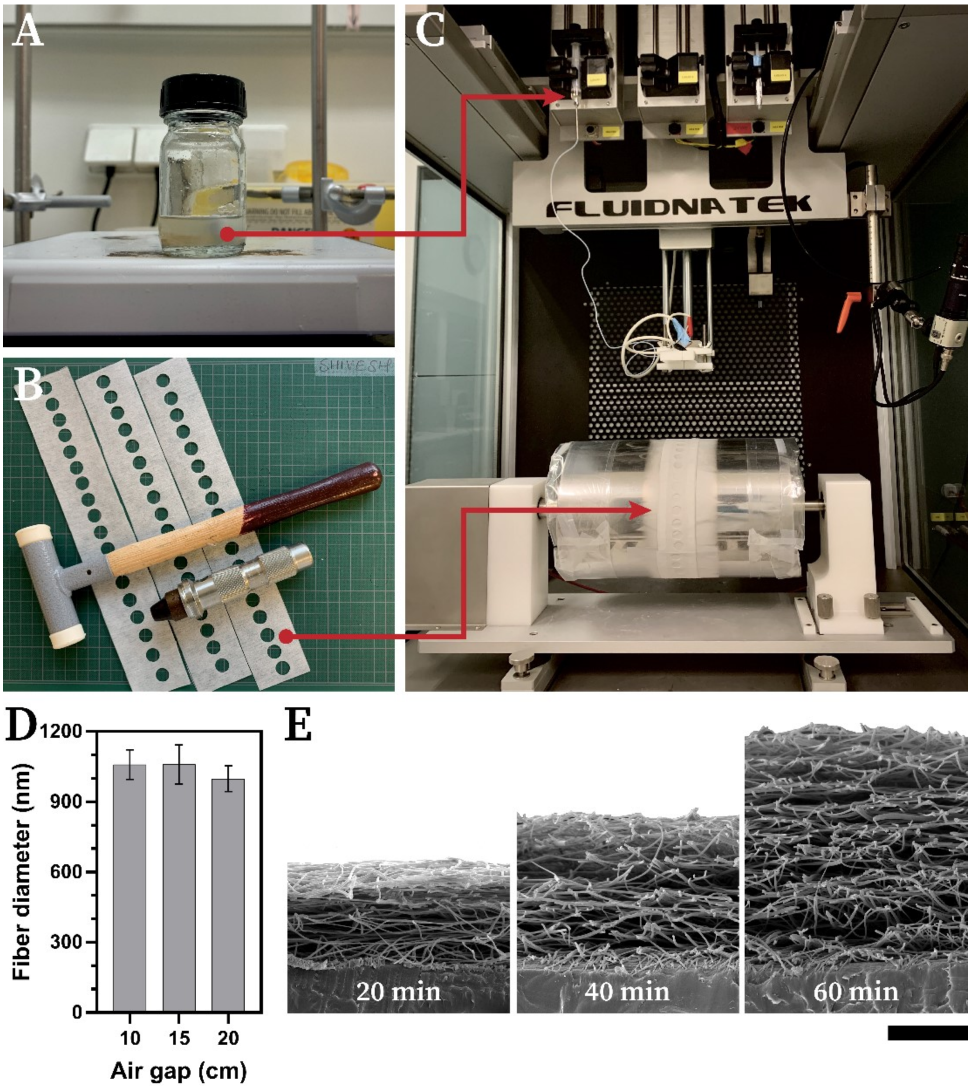


**Figure S1. Experimental setup for electrospinning. (A)** Preparation of PEOT/PBT copolymer solution in 70:30 chloroform and HFIP solvent mixture. **(B)** Preparation of collecting substrate, with punched Finishmat® 6691 (Lantor B.V., the Netherlands) over polypropylene (PP) sheets. **(C)** The electrospinning setup used for the fabrication of TM scaffolds. Arrows highlighting the placement of the polymer solution in syringe pump and PP sheet on the rotating mandrel. **(D)** Influence of the air gap between spinneret and collector on fiber diameter. **(E)** Fabrication of electrospun scaffolds with different thicknesses. Scale bar = 25 μm.

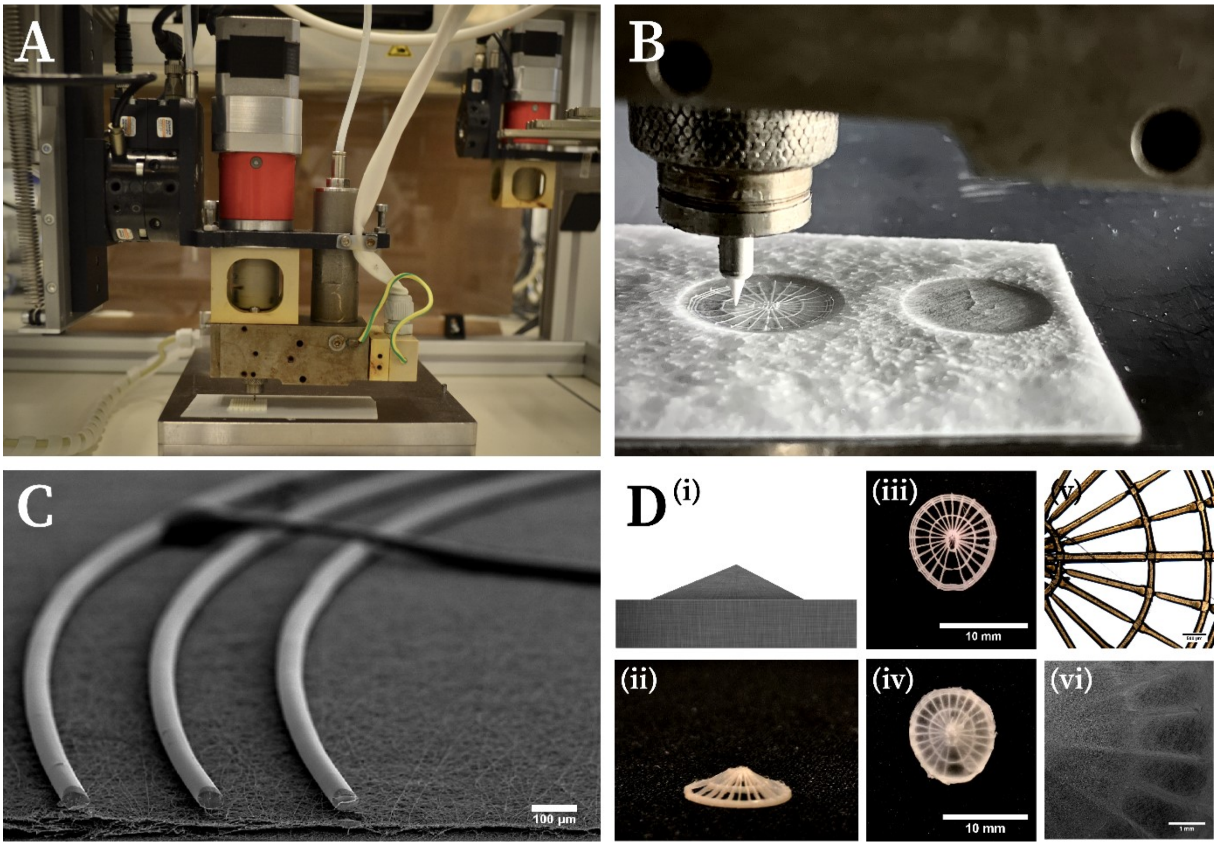


**Figure S2. Additive manufacturing of hierarchical TM constructs. (A)** The setup used for fused deposition modeling based 3D fiber deposition. **(B)** Deposition of a single layer of 50 μm filaments over electrospun meshes in pre-designed radial and circumferential patterns. **(C)** Scanning electron microscopy image highlighting the hierarchy in the fabricated hybrid scaffolds. **(D)** Manufacturing conical TM scaffolds: **(i)** 3D mold used as support for the fabrication, **(ii)** an example of conical TM scaffold, **(iii, iv)** top view of conical scaffolds before and after electrospinning, and **(v, vi)** microscopy images before and after electrospinning, respectively.

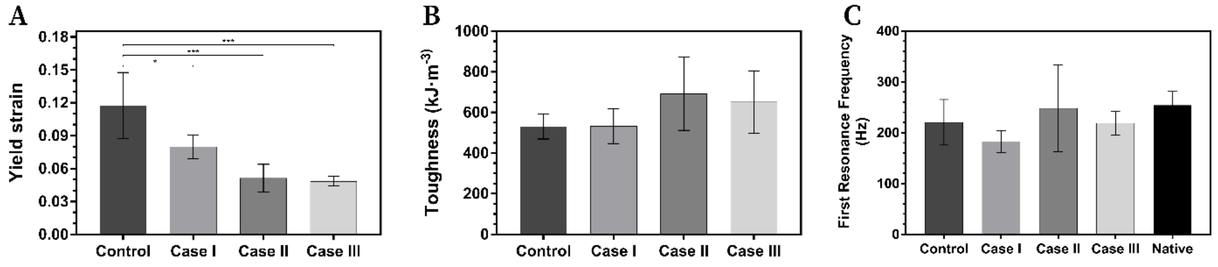


**Figure S3. Mechanical and acoustical characterization of the fabricated scaffolds. (A)** Calculated strain at the yield point. **(B)** Average toughness values summarized for all the cases. **(C)** Average first resonance frequencies for TM scaffolds fabricated with thicker electrospun mesh as compared to **Figure 4B**.

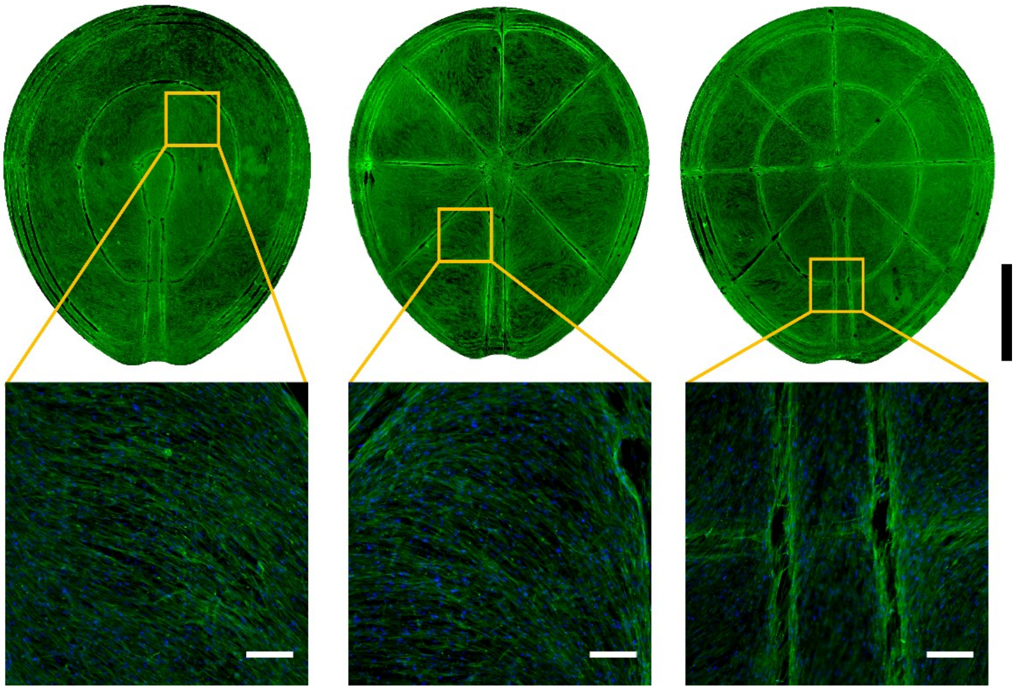

**Figure S4.** Phalloidin/DAPI staining at day 3 of human mesenchymal stromal cells seeded on the TM scaffolds; scale bar = 3 cm. Higher magnification images of select areas demonstrating cell alignment guided by the fabricated FDM filaments; scale bar = 100 μm.

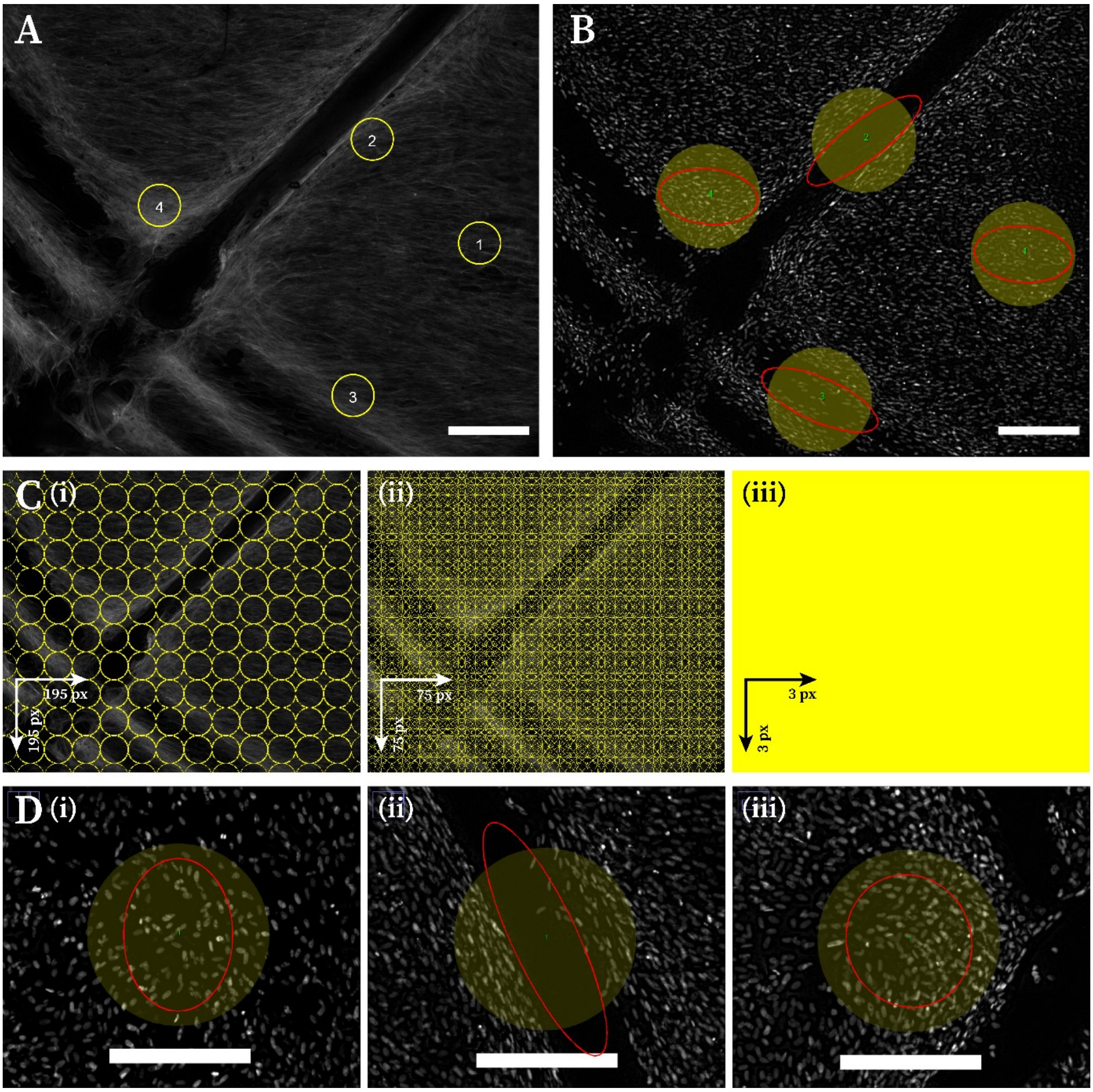


**Figure S5. Quantification of cell distribution and alignment. (A)** Demonstration of the four regions chosen for each image to compare the cell distribution: **(1)** no filament, **(2)** along radial filaments, **(3)** along circumferential filaments, and **(4)** junction between two filaments; scale bar = 250 µm. **(B)** Cell alignment quantification conducted in the same regions with OrientationJ plugin on Fiji. Gray-scale DAPI channel was used; scale bar = 250 µm. **(C)** Finding the ROI with the highest intensity for each image: **(i–iii)** demonstration of an increasing pixel shift for locating the most intense ROI; a 3 pixel shift resulting in generation of 641928 ROIs was used for the final normalization. **(D)** Examples highlighting the alignments obtained in the different regions. The yellow circle represents the ROI, and the red ellipse represents the extent of alignment: higher alignment (anisotropy) results in more elongated ellipse, as compared to completely rounded isotropic measurements. For example: **(i)** randomly scattered nuclei in a region without any filament – somewhat isotropic, **(ii)** aligned nuclei along the FDM filament – anisotropic, **(iii)** nuclei aligned in distinct orientations canceling out each other's effect at filament intersections – isotropic; scale bar = 250 μm.

**Table S1.** Summary of the mechanical characterization of each case with n = 4.

| Sample | ES thickness (μm) | Linear region (strain %) | | E (MPa) | $R^2$ value |
|---|---|---|---|---|---|
| | | Start | End | | |
| Control_1 | 8.46 | 1.50 | 9.00 | 5.65 | 0.98 |
| Control_2 | 9.77 | 1.50 | 10.50 | 4.72 | 0.98 |
| Control_3 | 9.77 | 3.00 | 16.00 | 3.42 | 0.99 |
| Control_4 | 9.77 | 2.50 | 11.50 | 3.72 | 0.98 |
| Case I_1 | 8.46 | 3.00 | 7.00 | 6.27 | 0.96 |
| Case I_2 | 9.77 | 4.50 | 8.00 | 5.80 | 0.97 |
| Case I_3 | 9.77 | 3.35 | 9.50 | 5.21 | 0.98 |
| Case I_4 | 8.46 | 3.35 | 7.50 | 7.14 | 0.97 |
| Case II_1 | 8.46 | 1.25 | 4.25 | 11.04 | 0.95 |
| Case II_2 | 9.77 | 2.20 | 4.95 | 7.95 | 0.95 |
| Case II_3 | 7.21 | 0.95 | 4.40 | 13.02 | 0.95 |
| Case II_4 | 8.46 | 4.30 | 6.95 | 7.41 | 0.95 |
| Case III_1 | 8.46 | 1.00 | 4.70 | 10.65 | 0.95 |
| Case III_2 | 8.46 | 1.50 | 4.50 | 12.31 | 0.97 |
| Case III_3 | 9.77 | 1.70 | 4.70 | 13.78 | 0.98 |
| Case III_4 | 7.21 | 2.50 | 5.50 | 12.34 | 0.95 |

### Python script for generating customized TM models:

```python
import math
import sys

sys.stdout = open("TM_v6.0_printer_Ceramic 70_on ES.nc","wt")

# ///////////////////////////////////////
# ////////// Parameters to set //////////
# ///////////////////////////////////////

inner_in = 0.50          # (1) Innermost ring
outer_in = 4.25          # (2) End of TM region, clamping begins (ratio of TM:clamping region)
ring = 4.75              # (3) End of clamping region, outermost ring
stretch = 1.50           # (4) Stretching of the ellipse
width_in = 2.0           # (5) Width between the rings in TM region
width_ot = 0.25          # (6) Width between the rings in clamping region
spokes = 24              # (7) Number of radial lines (density of spokes)
elevation = 2.197469     # (8) Depth of the cone


# /////////////////////////////////////////
# ////////// Printing parameters //////////
# /////////////////////////////////////////

temperature = 190        # Degree celcius
height = 0.07            # Height of the scaffold
feed = 157               # Feed-rate
zlayer_in = 0.05         # Layer height (in mm)
pressure = 250           # Pressure (in kPa)
screw = 5000             # Screw rotational speed (50 RPM)
tool = 11                # Tool number
needle = 9500            # Needle length (in um)

E4 = 0                   # Pull-back time (ms) [to activate the pull-back]
E7 = 0
E8 = 6000                # Pull-back speed (RPM)
E9 = 250                 # Pre-flow (ms)
E13 = 0                  # Post-flow (ms)
E21 = 0
E22 = 0


# /////////////////////////////////////////

inner = inner_in
outer = outer_in
zlayer = zlayer_in
elipse = ring+stretch

new_x_in = []            # creation of arrays for storing the coordinates
new_y_in = []            # creation of arrays for storing the coordinates

new_x_out = []
new_y_out = []

print ("G54")                    # coordinate system select
print (f"E21 {E21}")
print (f"E7 {E7}")
print (f"E111 {needle}")         # set needle length
print (f"E22 {E22}")
```

```python
print (f"T{tool} M06")          # tool change
print (f"E7 {E7}")
print (f"E0 {needle}")          # needlesensor command
print ("G00 Z30")
print ("G00 X0.0 Y0.0")
print (f"E6 {pressure}")


print ("G00 Z1.0")
while zlayer <= height:

    # ////////////////////////////////////////
    # / Construction of CIRCUMFERENTIAL lines /
    # ////////////////////////////////////////

    while inner <= outer:

        print (f"G00 X0.0 Y-{inner}")
        print (F"G01 Z{zlayer} F500")        # set z

        print ("E1 1")          # pressure ON
        print (f"E3 {screw}")   # set screw rotational speed
        print (f"E8 {E8}")
        print (f"E4 {E4}")
        print (f"E9 {E9}")
        print (f"E13 {E13}")
        print ("E2 1")          # screwing motor ON
        print (f"G03 X0.0 Y{inner} I0.0 J{inner} F{feed}")

        elipse = inner+stretch    # Stretching of the ellipse
        centre_x = 0
        centre_y = ((inner**2)-(elipse**2))/(2*inner)       # Centre of new circle = (0,centre_y)
        centre_1 = centre_y-inner
        centre_2 = -centre_y
        new_y = inner/3
        new_x = ((elipse**2)+(2*new_y*centre_y)-(new_y**2))**0.5

        print (f"G00 X0.0 Y{inner}")
        print (F"G01 Z{zlayer} F500")
        print (f"G03 X-{new_x} Y{new_y} I0.0 J{centre_1} F{feed}")

            if inner == inner_in:
                new_x_in.append(new_x)
                new_y_in.append(new_y)

            if inner > inner_in:
                print (f"G00 X-{new_x} Y{new_y}")
                print (f"G03 X-{new_x} Y{new_y/4} I{new_x-inner} J-{3*new_y/8} F{feed}")

                new_x_in.append(new_x)
                new_y_in.append(new_y/2)

                print (f"G00 X-{new_x} Y-{new_y/4}")
                print (f"G03 X-{new_x} Y-{new_y} I{new_x-inner} J-{3*new_y/8} F{feed}")


            print (f"G00 X-{new_x} Y-{new_y}")
            print (F"G01 Z{zlayer} F500")
```

```
            print (f"G03 X0.0 Y-{inner} I{new_x} J{centre_2+new_y} F{feed}")

            print ("E1 0")          # pressure OFF
            print ("E2 0")          # screwing motor OFF

            inner = inner+width_in     # Width between the rings in TM region

        while outer <= ring:

            print (f"G00 X0.0 Y-{outer}")
            print (F"G01 Z{zlayer} F500")

            print ("E1 1")          # pressure ON
            print (f"E3 {screw}")   # set screw rotational speed
            print (f"E8 {E8}")
            print (f"E4 {E4}")
            print (f"E9 {E9}")
            print (f"E13 {E13}")
            print ("E2 1")          # screwing motor ON
            print (f"G03 X0.0 Y{outer} I0.0 J{outer} F{feed}")

            elipse = outer+stretch
            centre_x = 0
            centre_y = ((outer**2)-(elipse**2))/(2*outer)
            centre_1 = centre_y-outer
            centre_2 = -centre_y
            new_y = outer/4
            new_x = ((elipse**2)+(2*new_y*centre_y)-(new_y**2))**0.5

            new_x_out.append(new_x)
            new_y_out.append(new_y/4)

            print (f"G00 X0.0 Y{outer}")
            print (F"G01 Z{zlayer} F500")
            print (f"G03 X-{new_x} Y{new_y} I0.0 J{centre_1} F{feed}")

            print (f"G00 X-{new_x} Y{new_y}")
            print (f"G03 X-{new_x} Y{new_y/4} I{new_x-outer*1.05} J-{3*new_y/8} F{feed}")

            print (f"G00 X-{new_x} Y{new_y/4}")
            print (f"G02 X-{new_x} Y-{new_y/4} I{new_x-outer*1.57} J-{new_y/4} F{feed}")    # The formula might change if the parameters are changed : 1.57 is chosen for the current version

            print (f"G00 X-{new_x} Y-{new_y/4}")
            print (f"G03 X-{new_x} Y-{new_y} I{new_x-outer*1.05} J-{3*new_y/8} F{feed}")

            print (f"G00 X-{new_x} Y-{new_y}")
            print (F"G01 Z{zlayer} F500")
            print (f"G03 X0.0 Y-{inner} I{new_x} J{centre_2+new_y} F{feed}")

            print ("E1 0")          # pressure OFF
            print ("E2 0")          # screwing motor OFF

            outer = outer+width_ot    # Width between the rings in clamping region

```

```python
        print (F"G01 Z{zlayer+1} F500")

        print (f"G00 X{ring} Y0.0")
        print (F"G01 Z{zlayer} F500")
        print ("E1 1")          # pressure ON
        print (f"E3 {screw}")   # set screw rotational speed
        print (f"E8 {E8}")
        print (f"E4 {E4}")
        print (f"E9 {E9}")
        print (f"E13 {E13}")
        print ("E2 1")          # screwing motor ON
        print (f"G01 X{inner_in} Y0.0 F{feed}")
        print ("E1 0")          # pressure OFF
        print ("E2 0")          # screwing motor OFF

        print (F"G01 Z{zlayer+1} F500")

        print (f"G00 X-{new_x_in[0]} Y{new_y_in[0]}")
        print (F"G01 Z{zlayer} F500")
        print ("E1 1")          # pressure ON
        print (f"E3 {screw}")   # set screw rotational speed
        print (f"E8 {E8}")
        print (f"E4 {E4}")
        print (f"E9 {E9}")
        print (f"E13 {E13}")
        print ("E2 1")          # screwing motor ON
        print (f"G01 X-{new_x_out[-1]} Y{new_y_out[2]} F{feed}")
        print ("E1 0")          # pressure OFF
        print ("E2 0")          # screwing motor OFF

        print (F"G01 Z{zlayer+1} F500")



        # ////////////////////////////////////////
        # ///// Construction of RADIAL lines //////
        # ////////////////////////////////////////

        theta_in = 360/spokes
        theta = theta_in

        radius = ((ring**2)+(elipse**2))/(2*ring)       # Radius of the outermost circle
        centre_y = ((ring**2)-(elipse**2))/(2*ring)     # Centre of the outermost circle = (0,centre_y). To calculate the intersection with y = x.tan(theta)

        elipse_in = inner_in + stretch
        radius_in = ((inner_in**2)+(elipse_in**2))/(2*inner_in)       # Radius of the innermost circle
        centre_y_in = ((inner_in**2)-(elipse_in**2))/(2*inner_in)     # Centre of the innermost circle = (0,centre_y_in). To calculate the intersection with y = x.tan(theta)

        while 0 <= theta < 90:
            rad = math.radians(theta)
            n = centre_y*math.tan(rad)+(radius**2+(radius*math.tan(rad))**2-centre_y**2)**0.5
            d = (1+(math.tan(rad))**2)
            x1 = n/d                            # Separated as numerator and denominator. Intersection of the circle and line
            if 0 < x1 < 10**(-6):
```

```
                x1 = 0
            x2 = ring*math.cos(rad)
            if 0 < x2 < 10**(-6):
                x2 = 0
            y1 = x1*math.tan(rad)
            if 0 < y1 < 10**(-6):
                y1 = 0
            y2 = ring*math.sin(rad)
            if 0 < y2 < 10**(-6):
                y2 = 0

            n_in = centre_y_in*math.tan(rad)+(radius_in**2+(radius_in*math.tan(rad))**2-centre_y_in**2)**0.5
            d_in = (1+(math.tan(rad))**2)
            x1_in = n_in/d_in                   # Separated as numerator and denominator. Intersection of the circle and line
            if 0 < x1_in < 10**(-6):
                x1_in = 0
            x2_in = inner_in*math.cos(rad)
            if 0 < x2_in < 10**(-6):
                x2_in = 0
            y1_in = x1_in*math.tan(rad)
            if 0 < y1_in < 10**(-6):
                y1_in = 0
            y2_in = inner_in*math.sin(rad)
            if 0 < y2_in < 10**(-6):
                y2_in = 0
            print (f"G00 X-{x1} Y{y1}")
            print (F"G01 Z{zlayer} F500")
            print ("E1 1")          # pressure ON
            print (f"E3 {screw}")   # set screw rotational speed
            print (f"E8 {E8}")
            print (f"E4 {E4}")
            print (f"E9 {E9}")
            print (f"E13 {E13}")
            print ("E2 1")          # screwing motor ON
            print (f"G01 X-{x1_in} Y{y1_in} F{feed}")
            print ("E1 0")          # pressure OFF
            print ("E2 0")          # screwing motor OFF
            print (F"G01 Z{zlayer+1} F500")

            print (f"G00 X{x2_in} Y{-y2_in}")
            print (F"G01 Z{zlayer} F500")
            print ("E1 1")          # pressure ON
            print (f"E3 {screw}")   # set screw rotational speed
            print (f"E8 {E8}")
            print (f"E4 {E4}")
            print (f"E9 {E9}")
            print (f"E13 {E13}")
            print ("E2 1")          # screwing motor ON
            print (f"G01 X{x2} Y-{y2} F{feed}")
            print ("E1 0")          # pressure OFF
            print ("E2 0")          # screwing motor OFF
            print (F"G01 Z{zlayer+1} F500")

            theta = theta+theta_in

            if theta==90:
                break

            rad = math.radians(theta)
```

```
            n = centre_y*math.tan(rad)+(radius**2+(radius*math.tan(rad))**2-centre_y**2)**0.5
            d = (1+(math.tan(rad))**2)
            x1 = n/d                             # Separated as numerator and denominator. Intersection of the circle and line
            if 0 < x1 < 10**(-6):
                x1 = 0
            x2 = ring*math.cos(rad)
            if 0 < x2 < 10**(-6):
                x2 = 0
            y1 = x1*math.tan(rad)
            if 0 < y1 < 10**(-6):
                y1 = 0
            y2 = ring*math.sin(rad)
            if 0 < y2 < 10**(-6):
                y2 = 0

            n_in = centre_y_in*math.tan(rad)+(radius_in**2+(radius_in*math.tan(rad))**2-centre_y_in**2)**0.5
            d_in = (1+(math.tan(rad))**2)
            x1_in = n_in/d_in                    # Separated as numerator and denominator. Intersection of the circle and line
            if 0 < x1_in < 10**(-6):
                x1_in = 0
            x2_in = inner_in*math.cos(rad)
            if 0 < x2_in < 10**(-6):
                x2_in = 0
            y1_in = x1_in*math.tan(rad)
            if 0 < y1_in < 10**(-6):
                y1_in = 0
            y2_in = inner_in*math.sin(rad)
            if 0 < y2_in < 10**(-6):
                y2_in = 0

            print (f"G00 X{x2} Y-{y2}")
            print (F"G01 Z{zlayer} F500")
            print ("E1 1")          # pressure ON
            print (f"E3 {screw}")   # set screw rotational speed
            print (f"E8 {E8}")
            print (f"E4 {E4}")
            print (f"E9 {E9}")
            print (f"E13 {E13}")
            print ("E2 1")          # screwing motor ON
            print (f"G01 X{x2_in} Y-{y2_in} F{feed}")
            print ("E1 0")          # pressure OFF
            print ("E2 0")          # screwing motor OFF
            print (F"G01 Z{zlayer+1} F500")

            print (f"G00 X-{x1_in} Y{y1_in}")
            print (F"G01 Z{zlayer} F500")
            print ("E1 1")          # pressure ON
            print (f"E3 {screw}")   # set screw rotational speed
            print (f"E8 {E8}")
            print (f"E4 {E4}")
            print (f"E9 {E9}")
            print (f"E13 {E13}")
            print ("E2 1")          # screwing motor ON
            print (f"G01 X-{x1} Y{y1} F{feed}")
            print ("E1 0")          # pressure OFF
            print ("E2 0")          # screwing motor OFF
            print (F"G01 Z{zlayer+1} F500")

```

```
            theta = theta+theta_in

        while theta==90:
            if theta_in % 2 == 0:
                print (f"G00 X0.0 Y{ring}")
                print (F"G01 Z{zlayer} F500")
                print ("E1 1")          # pressure ON
                print (f"E3 {screw}")   # set screw rotational speed
                print (f"E8 {E8}")
                print (f"E4 {E4}")
                print (f"E9 {E9}")
                print (f"E13 {E13}")
                print ("E2 1")          # screwing motor ON
                print (f"G01 X0.0 Y{inner_in} F{feed}")
                print ("E1 0")          # pressure OFF
                print ("E2 0")          # screwing motor OFF
                print (F"G01 Z{zlayer+1} F500")

                print (f"G00 X0.0 Y-{inner_in}")
                print (F"G01 Z{zlayer} F500")
                print ("E1 1")          # pressure ON
                print (f"E3 {screw}")   # set screw rotational speed
                print (f"E8 {E8}")
                print (f"E4 {E4}")
                print (f"E9 {E9}")
                print (f"E13 {E13}")
                print ("E2 1")          # screwing motor ON
                print (f"G01 X0.0 Y-{ring} F{feed}")
                print ("E1 0")          # pressure OFF
                print ("E2 0")          # screwing motor OFF
                print (F"G01 Z{zlayer+1} F500")

            else:
                print (f"G00 X0.0 Y-{ring}")
                print (F"G01 Z{zlayer} F500")
                print ("E1 1")          # pressure ON
                print (f"E3 {screw}")   # set screw rotational speed
                print (f"E8 {E8}")
                print (f"E4 {E4}")
                print (f"E9 {E9}")
                print (f"E13 {E13}")
                print ("E2 1")          # screwing motor ON
                print (f"G01 X0.0 Y-{inner_in} F{feed}")
                print ("E1 0")          # pressure OFF
                print ("E2 0")          # screwing motor OFF
                print (F"G01 Z{zlayer+1} F500")

                print (f"G00 X0.0 Y{inner_in}")
                print (F"G01 Z{zlayer} F500")
                print ("E1 1")          # pressure ON
                print (f"E3 {screw}")   # set screw rotational speed
                print (f"E8 {E8}")
                print (f"E4 {E4}")
                print (f"E9 {E9}")
                print (f"E13 {E13}")
                print ("E2 1")          # screwing motor ON
                print (f"G01 X0.0 Y{ring} F{feed}")
                print ("E1 0")          # pressure OFF
                print ("E2 0")          # screwing motor OFF
                print (F"G01 Z{zlayer+1} F500")

            theta = theta+theta_in
```

```
        while 90 < theta < 180:
            rad = math.radians(theta)
            n = -centre_y*math.tan(rad)+(radius**2+(radius*math.tan(rad))**2-centre_y**2)**0.5
            d = (1+(math.tan(rad))**2)
            x1 = n/d                                   # Separated as numerator and denominator. Intersection of the circle and line
            if 0 < x1 < 10**(-6):
                x1 = 0
            x2 = -ring*math.cos(rad)
            if 0 < x2 < 10**(-6):
                x2 = 0
            y1 = x1*math.tan(rad)
            if 0 < y1 < 10**(-6):
                y1 = 0
            y2 = ring*math.sin(rad)
            if 0 < y2 < 10**(-6):
                y2 = 0

            n_in = -centre_y_in*math.tan(rad)+(radius_in**2+(radius_in*math.tan(rad))**2-centre_y_in**2)**0.5
            d_in = (1+(math.tan(rad))**2)
            x1_in = n_in/d_in                          # Separated as numerator and denominator. Intersection of the circle and line
            if 0 < x1_in < 10**(-6):
                x1_in = 0
            x2_in = -inner_in*math.cos(rad)
            if 0 < x2_in < 10**(-6):
                x2_in = 0
            y1_in = x1_in*math.tan(rad)
            if 0 < y1_in < 10**(-6):
                y1_in = 0
            y2_in = inner_in*math.sin(rad)
            if 0 < y2_in < 10**(-6):
                y2_in = 0

            print (f"G00 X-{x1} Y{y1}")
            print (F"G01 Z{zlayer} F500")
            print ("E1 1")          # pressure ON
            print (f"E3 {screw}")   # set screw rotational speed
            print (f"E8 {E8}")
            print (f"E4 {E4}")
            print (f"E9 {E9}")
            print (f"E13 {E13}")
            print ("E2 1")          # screwing motor ON
            print (f"G01 X-{x1_in} Y{y1_in} F{feed}")
            print ("E1 0")          # pressure OFF
            print ("E2 0")          # screwing motor OFF
            print (F"G01 Z{zlayer+1} F500")

            print (f"G00 X{x2_in} Y{y2_in}")
            print (F"G01 Z{zlayer} F500")
            print ("E1 1")          # pressure ON
            print (f"E3 {screw}")   # set screw rotational speed
            print (f"E8 {E8}")
            print (f"E4 {E4}")
            print (f"E9 {E9}")
            print (f"E13 {E13}")
            print ("E2 1")          # screwing motor ON
            print (f"G01 X{x2} Y{y2} F{feed}")
```

```
        print ("E1 0")          # pressure OFF
        print ("E2 0")          # screwing motor OFF
        print (F"G01 Z{zlayer+1} F500")

        theta = theta+theta_in

        if theta==180:
            break

        rad = math.radians(theta)
        n = -centre_y*math.tan(rad)+(radius**2+(radius*math.tan(rad))**2-centre_y**2)**0.5
        d = (1+(math.tan(rad))**2)
        x1 = n/d                            # Separated as numerator and denominator. Intersection of the circle and line
        if 0 < x1 < 10**(-6):
            x1 = 0
        x2 = ring*math.cos(rad)
        if 0 < x2 < 10**(-6):
            x2 = 0
        y1 = x1*math.tan(rad)
        if 0 < y1 < 10**(-6):
            y1 = 0
        y2 = ring*math.sin(rad)
        if 0 < y2 < 10**(-6):
            y2 = 0

        n_in = -centre_y_in*math.tan(rad)+(radius_in**2+(radius_in*math.tan(rad))**2-centre_y_in**2)**0.5
        d_in = (1+(math.tan(rad))**2)
        x1_in = n_in/d_in                   # Separated as numerator and denominator. Intersection of the circle and line
        if 0 < x1_in < 10**(-6):
            x1_in = 0
        x2_in = inner_in*math.cos(rad)
        if 0 < x2_in < 10**(-6):
            x2_in = 0
        y1_in = x1_in*math.tan(rad)
        if 0 < y1_in < 10**(-6):
            y1_in = 0
        y2_in = inner_in*math.sin(rad)
        if 0 < y2_in < 10**(-6):
            y2_in = 0

        if x2 < 0:
            x2_var = -x2
            print (f"G00 X{x2_var} Y{y2}")
        else:
            print (f"G00 X-{x2} Y{y2}")
        print (F"G01 Z{zlayer} F500")
        print ("E1 1")          # pressure ON
        print (f"E3 {screw}")   # set screw rotational speed
        print (f"E8 {E8}")
        print (f"E4 {E4}")
        print (f"E9 {E9}")
        print (f"E13 {E13}")
        print ("E2 1")          # screwing motor ON
        if x2_in < 0:
            x2_in_var = -x2_in
            print (f"G01 X{x2_in_var} Y{y2_in} F{feed}")
        else:
```

```
                print (f"G01 X-{x2_in} Y{y2_in} F{feed}")
            print ("E1 0")          # pressure OFF
            print ("E2 0")          # screwing motor OFF
            print (F"G01 Z{zlayer+1} F500")

            print (f"G00 X-{x1_in} Y{y1_in}")
            print (F"G01 Z{zlayer} F500")
            print ("E1 1")          # pressure ON
            print (f"E3 {screw}")   # set screw rotational speed
            print (f"E8 {E8}")
            print (f"E4 {E4}")
            print (f"E9 {E9}")
            print (f"E13 {E13}")
            print ("E2 1")          # screwing motor ON
            print (f"G01 X-{x1} Y{y1} F{feed}")
            print ("E1 0")          # pressure OFF
            print ("E2 0")          # screwing motor OFF
            print (F"G01 Z{zlayer+1} F500")

            theta = theta+theta_in

        print (f"G00 X-{new_x_in[0]} Y-{new_y_in[0]}")
        print (F"G01 Z{zlayer} F500")
        print ("E1 1")          # pressure ON
        print (f"E3 {screw}")   # set screw rotational speed
        print (f"E8 {E8}")
        print (f"E4 {E4}")
        print (f"E9 {E9}")
        print (f"E13 {E13}")
        print ("E2 1")          # screwing motor ON
        print (f"G01 X-{new_x_out[-1]} Y-{new_y_out[2]} F{feed}")
        print ("E1 0")          # pressure OFF
        print ("E2 0")          # screwing motor OFF

        print (F"G01 Z{zlayer+1} F500")

        zlayer = zlayer+zlayer_in

        inner = inner_in            # reset the value for next layer
        outer = outer_in
        theta = theta_in

    sys.stdout.close()
```
